# Cholesterol and high concentration serum cooperatively enhance canine CAR T cell transduction and expansion

**DOI:** 10.64898/2026.11.25.769304

**Authors:** Treyvon Davis, Micah Mallory, Mark Ogden, Christopher Moody, Hayden Sandt, Carla O’Reilly, Stephen Gottschalk, Christopher DeRenzo, Christopher Mariani, Paul Hess, Jennifer Holmes, Steven Suter, Yevgeny Brudno

## Abstract

Chimeric antigen receptor (CAR) T cell immunotherapy is emerging as a promising adjunct treatment option for refractory canine cancers. Unfortunately, clinical trials thus far have failed to meaningfully reduce tumor burden, highlighting the need for improved CAR T cell manufacturing strategies. Effective autologous therapy requires highly viable, efficiently transduced T cells capable of robust expansion, sustained CAR expression, and long-term persistence. Although these parameters have been extensively investigated in humans, comparable optimization in canine systems remains limited. Here, we evaluated culture conditions to improve canine T cell expansion, memory differentiation, and retroviral transduction efficiency. We demonstrated IL-2 supplementation supported superior cellular expansion and preservation of less-differentiated memory phenotypes compared to IL-7 with IL-15. Additionally, we found that increasing cell density during shorter stimulation periods improved expansion while also driving better T cell transduction rates. We discovered that the addition of cholesterol and high-concentration fetal bovine serum (FBS) to media during the transduction period reliably and markedly enhanced transduction efficiency following a single round of retroviral transduction, even at elevated temperatures that would otherwise impair transduction efficiency. This enhancement was observed with both GALV- and RD114-pseudotyped retroviruses and was accompanied by a marked improvement in cellular expansion without driving terminal differentiation of T cell subsets that are prone to exhaustion *in vivo*. Collectively, these findings identify practical strategies to improve canine CAR T-cell manufacturing and advance the development of clinically viable CAR T cells for canine cancer treatment.

## Introduction

Chimeric antigen receptor (CAR) T cell therapy has revolutionized the treatment of human malignancies, yet its development in companion animals lags behind advances in human medicine [1]. Closing this gap has dual relevance. First, canine cancers are increasingly valuable comparative models whose study can accelerate progress for human cellular immunotherapy. Second, dogs are patients in their own right who stand to benefit from novel therapeutic alternatives to conventional radiation, chemotherapeutic, or surgical treatment strategies.

Naturally occurring cancers in dogs are increasingly recognized as valuable comparative models of human malignancies, arising spontaneously in immunocompetent hosts and exhibiting both tumor heterogeneity and microenvironmental complexities that are difficult to replicate in induced and xenograft mouse models. The translational value of canine cancer is further supported by similarities between aggressive cancers in dogs and humans. Canine and human malignancies can share patterns of biological behavior, including local invasion and metastatic potential, have similar genetic origins, and respond to similar treatment protocols [2–6]. For some cancers, the substantially higher incidence in dogs also creates opportunities to study diseases that are relatively rare in humans. Osteosarcoma, for example, has a more than 14-fold higher incidence in dogs in the United States, with more than 8,000 canine cases seen annually [7,8]. Clinical evaluation of CAR T cell therapy in dogs may also provide valuable insights into lymphodepletion requirements, treatment-related toxicities, and anti-CAR immune responses, which are important considerations in human trials but can be difficult to adequately model in rodents [9–13].

Beyond their role as models, dogs with aggressive or refractory cancers have few effective treatment options. Surgery, radiation therapy, and chemotherapy remain the standard of care for many aggressive tumors, including osteosarcoma, visceral hemangiosarcoma, and glioma. Unfortunately, with conventional therapy, median survival of patients with these cancers remains below 12, 6, and 3 months respectively [14–16]. Even in cancers that are responsive to chemotherapy, such as canine multicentric lymphoma (70-90% remission), disease recurrence is almost certain and is increasingly refractory to treatment, limiting median survival time to 8-12 months [17]. These therapeutic limitations underscore the need for alternative treatment strategies, including targeted immunotherapies.

CAR T cells have shown promise in veterinary medicine. Clinical trials utilizing CD20-specific CAR T cells in canine patients with diffuse large B cell lymphoma (DLBCL) resulted in reduced numbers of CD20-expressing B cells in peripheral blood and malignant lymph nodes [18,19]. These trials have made promising advancements in canine CAR T cell research, but incomplete tumor elimination and lack of CAR T cell persistence demonstrated that more work needs to be done to produce CAR T cell infusion products in dogs that provide robust anti-tumor effects while maintaining durability and longevity.

Effective autologous CAR T cell therapy depends on producing highly viable, efficiently transduced T cells capable of sustained CAR expression, robust expansion, and long-term persistence following adoptive transfer. While considerable progress has been made toward these goals in humans, the veterinary field has yet to systematically optimize the manufacturing parameters of a successful autologous product in dogs. CAR T cells capable of high survival and expansion rates must first receive adequate stimulation. As such, we previously optimized stimulatory antibody choices and presentation strategies to induce sustained canine T cell activation and expansion [20]. In addition, maintaining memory CAR T cell populations is likely crucial for durability in vivo. In human CAR T cell clinical trials, patients whose cells contained higher proportions of less differentiated memory T cells, namely central memory (T_CM_) and stem cell memory (T_SCM_) T cells had better outcomes [21,22]. Effector (T_EFF_) T cells are critical for tumor cell engagement, however these terminally differentiated cells, are prone to exhaustion and need to be replenished by less differentiated cell precursors for enduring tumor destruction [22–24]. Recombinant human (rh)IL-2 is well known to drive excellent expansion of activated canine T cells and may outperform canine IL-2 [8,20,25–28], but limited attention has been given to its effect on T cell memory in dogs. IL-7 and IL-15 have been used to successfully preserve less differentiated memory phenotypes in human CAR T cells [29], but are rarely explored in canine studies.

Achieving reliably high transduction efficiency is crucial for generating clinically viable CAR T cell products, but this vital manufacturing step has received little systematic attention in canine systems. In fact, most publications attempting canine CAR T cell engineering fail to report transduction rates consistently above 30-40%. With the inherent difficulty in successful transduction, groups often implement strategies to boost transduction rates, including repeated rounds of viral transduction or temporary co-culture with target cells to enrich for CAR T cells [19,26–28,30]. While these strategies do improve transduction, they ultimately raise cost, increase culture time, and, in the case of co-culture, may drive premature T cell exhaustion. In our previous work, we established that plate-bound antibody stimulation resulted in transduction rates consistently above 50% with a single round of transduction [20]. These experiments, however, used a GFP retrovirus and not the more clinically relevant CAR retrovirus.

In this work, we build a framework for optimized canine CAR T cell immunotherapy, characterized by high T cell fitness, robust expansion of less differentiated memory subsets, and high transduction rates with sustained CAR expression. We compare the effects of recombinant human IL-2, IL-7, and IL-15 in culture and demonstrate IL-2 supplementation promotes excellent viability and drives better expansion with improved memory T cell profiles. We also discovered that cholesterol in combination with high concentration fetal bovine serum (FBS) dramatically boosts transduction rates in canine T cells, resulting in single-round transduction rates consistently above 70%, and frequently exceeding 80%. We highlight that this combination boosts transduction rates across multiple retroviral envelopes while dramatically improving expansion profiles without driving terminal T_EFF_ differentiation. Collectively, these optimizations address gene transfer and expansion bottlenecks that constrain canine CAR T cell manufacturing, yielding a highly transduced and less-differentiated canine CAR T cell product that has been associated with stronger and more durable responses in human patients.

## Materials and methods

### PBMC isolation

For canine PBMCs, blood samples were collected from healthy canine donors by the North Carolina State University Lab Animal Resources veterinary team. Relevant donor information is provided in S1 Table. For processing, 15 mL of Lymphoprep (Accurate Chemical) was added to 50 mL conical tubes. Whole blood was diluted 1:1 with 1X Dulbecco’s phosphate-buffered saline (DPBS) (Gibco), and 25 mL of the diluted blood was carefully added to the Lymphoprep layer. Samples were centrifuged at 400 x g for 40 minutes using the lowest acceleration setting and no brake. The peripheral blood mononuclear cell (PBMC) layer was then collected, transferred to clean 50 mL tubes, and washed twice with DPBS and centrifugation. The resulting PBMC pellet was resuspended in cold freezing media composed of 50% fetal bovine serum (FBS), 40% complete canine T cell media, and 10% DMSO (Sigma). Aliquots were frozen at −80°C before being transferred to liquid nitrogen.

For human PBMCs, donor blood was purchased from Gulf Coast Blood Center. PBMC isolation and cold storage were performed in a similar fashion to canine cells. Human PBMC freezing media was composed of 50% fetal bovine serum (FBS), 40% complete human T cell media, and 10% DMSO. *Macaca nemestrina* non-human primate (NHP) PBMCs were obtained from blood samples collected from donors (Washington National Primate Research Center) through Percoll density gradient separation. In short, the cells were initially spun down (1400 x g for 15 minutes) and excess plasma removed, diluted in 1 X HBSS with EDTA (5.39 mM), added to the top of a warmed Percoll solution and spun again (1000 x g for 30 minutes with brakes off). Then, the band of white blood cells was collected and washed. NHP PBMCs were frozen in 90% complete NHP T cell media with 10% DMSO.

### Canine T cell culture and activation

Complete canine T cell media (cTCM) was prepared with Advanced RPMI 1640 (Gibco) supplemented with 10 mmol/L HEPES (pH 7.3, ThermoScientific), 2 mmol/L L-Glutamine (GlutaMAX, Gibco), 100 U/mL Penicillin and 100 μg/mL Streptomycin (Gibco), and 10% FBS (Corning 35-015-CV). Serum free media (SFM) was prepared as cTCM, but without any FBS. Phosphate-free media was prepared using RPMI 1640 without L-Glutamine and Phosphate (MP Biomedicals) supplemented with 10 mmol/L HEPES, 1 mmol/L Sodium Pyruvate (Gibco), 2 mmol/L L-Glutamine, 100 U/mL Penicillin and 100 μg/mL Streptomycin, with or without 3% FBS. Prior to activation, T cells were thawed and allowed to rest in cTCM overnight in a humidified incubator at 38.8°C, 5% CO_2_.

T cells were activated using a slightly modified protocol described previously [20]. Briefly, for T cell activation, stimulatory anti-canine CD3 antibody (clone CA17.6F9, purchased from UC Davis Leukocyte Antigen Biology Laboratory) and anti-canine CD28 antibody (clone 5B8, purified into PBS from hybridoma generously gifted by Brian Hayes, Fred Hutchinson Cancer Research Center) were first bound to nontreated polystyrene 24-well plates (Genesee) as follows: Antibody solution was prepared in distilled water with αCD3 at 2 μg/mL and αCD28 at 4 μg/mL. 500 μL was added to each well. The plate was placed in 4°C overnight. Plates were then removed and the antibody solution was aspirated from the wells. 2 mL cTCM containing 10% FBS was added to each well and incubated for 15–30 minutes at 38.8°C as a blocking step [31] and to remove free antibody prior to addition of PBMCs. In initial experiments, 1 x 10^6^ PBMCs in 2 mL of cTCM supplemented with rhIL-2 (Peprotech) (200 IU/mL for all experiments unless otherwise specified) were added to each well containing bound antibody and returned to the incubator. Cells were stimulated for 2-5 days before being removed from antibody-coated plates, and expanded at 38.8°C in cTCM supplemented with IL-2.

In subsequent experiments, we increased initial PBMC seeding density in antibody-coated plates to enhance cellular expansion. For 2-day stimulation groups, seeding density was doubled from 1 x 10^6^ to 2 x 10^6^ cells/well (5.26 x 10^5^ cells/cm^2^ to 1.05 x10^6^ cells/cm^2^). For 3-day stimulation groups, seeding density was increased to 1.5 x 10^6^ cells/well (7.89 x10^5^ cells/cm^2^) to accommodate the additional day of proliferation while minimizing overcrowding and preserving viability.

### Cytokine Assays

rhIL-2 (Peprotech), rhIL-7 (Peprotech), and rhIL-15 (Peprotech) were evaluated in the context of T cell expansion, viability, and phenotype differentiation at different concentrations. To evaluate cytokine effects in the presence of stimulatory antibodies, 1 x 10^6^ PBMCs from five healthy canine donors were added to antibody-coated plates for 5 days in cTCM media supplemented with IL-2 at 100, 200, or 500 IU/mL or with IL-7+IL-15 at 5+2.5, 10+5, or 20+10 ng/mL. Media containing no cytokines was used as a negative control. On days 3 and 5, cells were evaluated for CD4/CD8 and memory T cell subsets by flow cytometry. On day 5, cell counts were performed using CellDrop (DeNovix).

In a separate experiment run concurrently, PBMCs activated in like fashion were removed from stimulation after 3 days. To determine cytokine effects during the expansion period, 10^6^ cells from each donor/condition were then transferred to T25 flasks (Corning) containing cTCM and expanded for 4 days in their respective cytokines at maintained concentrations. On day 4, cell counts were performed using CellDrop, and phenotype and viability analysis were performed using flow cytometry.

Importantly for IL-2, we used “international units” (IU) as established by WHO international standards, which are different from the term “units” used by PeproTech as defined by biological activity (WHO: 2180 IU/μg; Peprotech: ED50 as determined by the dose-dependent stimulation of murine CTLL-2 cells is ≤ 0.1 ng/ml, corresponding to a specific activity of ≥ 1 x 10^7^ units/mg, according to datasheet information).

### Canine T cell transduction assays

For RetroNectin-mediated transduction, 7 μL RetroNectin (TaKaRa T100A) in 1 mL DPBS was added to each well of nontreated 24-well plates the day prior to transduction, and incubated at 4°C overnight. The following day, RetroNectin was aspirated and plates were blocked with cTCM containing 10% FBS. cTCM was removed prior to the addition of GALV-pseudotyped B7-H3 CAR retrovirus, RD114-pseudotyped B7-H3 CAR retrovirus, or RD114-pseudotyped GFP retrovirus. Plates were centrifuged at 2000 x g for 90 minutes at 32°C. Viral supernatant was aspirated and 10^6^ activated T cell-enriched PBMCs were added to the wells of the RetroNectin/virus coated plates in 2 mL IL-2 supplemented cTCM. Plates were centrifuged at 1000 x g for 10 minutes and placed in a 37°C incubator overnight prior to transferring to 38.8°C incubator for expansion, unless otherwise specified. Variable incubator temperatures were used because our retrovirus showed reduced infectivity at higher temperatures, but canine T cells respond better to stimulation at 38.8°C and also have better expansion rates [20,32]. All samples were limited to a single round of transduction. Cells were expanded for 7 days with fresh IL-2 enriched media provided every 3 days. CAR expression was evaluated by flow cytometry to determine transduction efficiency.

### Human and NHP T cell culture, activation, and transduction

Human complete T cell media (hTCM) was prepared using 1:1 v/v Click’s Medium (Fujifilm Irvine Scientific) and HyClone RPMI 1640 (Cytiva) supplemented with 10% FBS, 2 mmol/L GlutaMAX, 100 U/mL Penicillin, and 100 μg/mL Streptomycin. For T cell activation, human PBMCs were thawed one day prior to activation and rested overnight in T cell media. Non-treated 24 well tissue culture plates were coated with 0.5 μg αCD3 antibody (Miltenyi Biotec, clone OKT3) and 0.5 μg αCD28 antibody (BD Biosciences, clone CD28.2) in 500 μL water, and left overnight at 4°C. The next day, wells were washed and blocked with 500 μL T cell media for 30 minutes at 37°C, after which 1×10^6^ PBMCs in 2 mL (hTCM) were added to each well and cultured for 48 hours at 37°C. After 48 hours, 250 IU/mL rhIL2 was added to each well and cultured for an additional 24 hours. Cells were then collected off the coated plates and immediately used for viral transduction experiments in like manner to canine cells. Transduction efficiency was evaluated on day 5.

NHP complete T cell media (nhpTCM) was prepared using X-VIVO Media (Lonza) supplemented with 10% FBS, 100 U/mL Penicillin and 100 μg/mL Streptomycin, and 50 μM β-mercaptoethanol. For NHP T cell activation, non-treated 24 well tissue culture plates were coated with 0.5 μg αCD3 antibody (BD Biosciences, clone SP34-2) and 0.5 μg αCD28 antibody (BD Biosciences, clone CD28.2) in 500 μL water, and left overnight at 4°C. The next day, wells were washed and blocked with 500 μL T cell media for 30 minutes at 37°C, after which 1×10^6^ PBMCs in 2 mL (nhpTCM) were added to each well and cultured for 48 hours at 37°C. After 48 hours, 250 IU/mL rhIL2 was added to each well and cultured for 24 hours. Cells were then collected off the coated plates and immediately used for viral transduction experiments in like manner to canine cells. Transduction efficiency was evaluated on day 3.

### GALV-retrovirus transduction after phosphate and serum deprivation

For phosphate deprivation transduction experiments, PBMCs from four healthy canine donors were activated for 2 days as described above. At 16 hours or 6 hours prior to transduction, cells were collected from antibody coated plates and centrifuged at 400xg for 5 minutes. 1 mL phosphate-free media (3% FBS) containing 200 IU/mL IL-2 was placed in antibody-coated plates during centrifugation to prevent them from drying out. After centrifugation, supernatant was removed and cells were resuspended in IL-2 enriched phosphate free media (3% FBS) and immediately returned to antibody coated plates. A negative control group did not undergo phosphate deprivation and was maintained in IL-2 enriched cTCM. At transduction, all cells were returned to IL-2 enriched standard cTCM (10% FBS) and transduced as described above with GALV-pseudotyped B7-H3 retrovirus. The following day after transduction, cells from all groups were transferred directly to T25 flasks containing 10 mL standard IL-2 enriched cTCM (10% FBS), with fresh cytokine-containing media added after 3 days. Transduction efficiency was evaluated at day 4 and 7 by flow cytometry.

For phosphate and serum deprivation experiments, activation and transduction were performed in a similar fashion with the following changes: 12 hours and 6 hours prior to transduction, media was replaced with IL-2 enriched phosphate-free media without FBS. No changes were made to the negative control group. For an additional group, 6 hours prior to transduction media was replaced with IL-2 enriched phosphate-free media without FBS, and cells were maintained in this media overnight during transduction. For all other groups, cells were returned to IL-2 enriched standard cTCM during the transduction period. The following day after transduction, cells from all groups were transferred directly to T25 flasks containing 10 mL standard IL-2 enriched cTCM (10% FBS). On Day 4, 1 mL from each flask was collected to assess transduction and the remaining cells were transferred to T75 flasks with fresh IL-2 enriched media for further expansion. Cell counts using CellDrop and transduction efficiency by flow cytometry were determined on day 7.

### Transduction after supplementation with high concentration FBS, cholesterol, or BSA

For experiments evaluating transduction at different concentrations of FBS, PBMCs from four healthy canine donors were activated for 2 days as described above. At the time of transduction, 10^6^ cells were transferred to 2 mL IL-2 enriched SFM or cTCM with different concentrations of FBS (10%, 20%, 30%, 40%, or 50%) and transduced with GALV retrovirus as described above. The following day, the 2 mL cell suspension was transferred directly to T25 flasks containing 10 mL standard IL-2 enriched cTCM (10% FBS). On day 4, cells were transferred to T75 flasks with fresh IL-2 enriched media for further expansion. On day 7, transduction efficiency was determined by flow cytometry. Median fluorescence intensity (MFI) was also evaluated as an indirect indicator of transgene expression levels.

For experiments comparing the effects of FBS, cholesterol, and bovine serum albumin (BSA) on transduction, activation and transduction were performed in a similar fashion with the following changes: At the time of transduction, 10^6^ cells were transferred to 2 mL IL-2 enriched media with different concentrations of FBS or cholesterol (250X Cholesterol Lipid Concentrate, Gibco 12531-018; exact molar concentration not disclosed by the manufacturer), BSA (BSA Fraction V, Gibco 15260037), FBS+cholesterol, or FBS+BSA. In all instances, BSA concentration was equivalent to BSA in a 30% FBS solution according to the Corning certificate of analysis for the particular FBS lot number. Cells were transduced with GALV or RD114-pseudotyped retrovirus as described above. The following day, the 2 mL cell suspension was transferred directly to T25 flasks containing 10 mL standard IL-2 enriched cTCM (10% FBS).

On day 4, cell counts were performed using CellDrop before being transferred to T75 flasks with fresh IL-2 enriched media for further expansion. On day 7 cell counts were performed and transduction efficiency and MFI was evaluated by flow cytometry. In subsequent experiments evaluating the impact of FBS+cholesterol on memory and CD4/CD8 phenotypes, phenotype analysis was performed on day 4 of expansion by flow cytometry.

For experiments evaluating the effects of higher culture temperatures during the transduction period, 10^6^ activated T cell-enriched PBMCs from four healthy canine donors were transduced either at 37°C or at 38.8°C overnight. The following day, cells were transferred directly to IL-2 enriched cTCM and expanded at 38.8°C. On day 7 post-transduction, transduction efficiency was evaluated by flow cytometry and the relative reduction in transduction was evaluated for each individual donor from each group using the following equation:

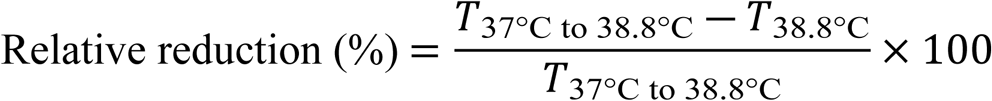

Experiments evaluating the effects of canine plasma (CP) on transduction used sterile-filtered plasma that was collected at the time of PBMC isolation and frozen. Plasma was thawed before use. Canine cells from two healthy donors were activated as above and 10^6^ cells were transferred to 2 mL IL-2 enriched media containing 0% or 10% FBS, or cholesterol with or without 30% FBS or 10% CP and transduced with GALV retrovirus as described above. Higher concentrations of plasma could not be evaluated as significant fibrin clot formation was observed after addition to media which interfered with T cell viability. The following day, the 2 mL cell suspension was transferred directly to 10 mL standard IL-2 enriched cTCM for expansion. On day 7 post-transduction, transduction efficiency was evaluated by flow cytometry.

### Flow cytometry

For viability assessment, cell suspensions were collected and washed once with DPBS and centrifugation at 400xg for 5 minutes. Cells were stained with LIVE/DEAD Near IR Fixable Dead Cell Stain Kits (Invitrogen) or Live-or-Dye 405/452 Fixable Viability Staining Kits (Biotium) prepared at 1:1000 dilution in 1X DPBS and incubated in the dark at 4°C for 30 minutes. Cells were washed in DPBS and again in Fetal Bovine Serum Stain Buffer (BD) before adding labeling antibodies.

To evaluate T cell subsets, anti-canine CD5 (clone YKIX322.3)-PerCP-eFluor 710 (Invitrogen, eBioscience), anti-canine CD8a (clone YCATE55.9)-Super Bright 436 or APC (Invitrogen, eBioscience), anti-canine CD4 (clone YKIX302.9)-Super Bright 436 or APC (Invitrogen, eBioscience), anti-human CD62L (clone FMC46)-FITC (Bio-Rad, validated in dogs [33]), and anti-canine CD45RA (clone CA21.4B3-biotinylated, purchased from UC Davis Leukocyte Antigen Biology Laboratory) were added to cell suspension and incubated in the dark at 4°C for 30 minutes. After washing twice with stain buffer, cells were finally stained with Streptavidin-PE (Invitrogen, eBioscience) specific for biotinylated antibody. In some instances, anti-canine CD25 (clone P4A10)-Super Bright 600 (Invitrogen, eBioscience) was used to label activated T cells. To determine transduction efficiency, CAR protein was labeled with αG4S linker (clone E7O2V)-PE (Cell Signaling Technology) and incubated in the dark at 4°C for 30 minutes.

All antibody and streptavidin solutions to label cell surface markers were prepared in stain buffer at the appropriate antibody dilution based on titration studies using flow cytometry. After antibody labeling, cells were fixed using 250 μL Cytofix Fixation Buffer (BD), except GFP-transduced cells, which were not fixed. Cells were then washed once and resuspended in stain buffer for flow cytometry.

Human and NHP cells were prepared in a similar fashion to canine cells. Human T cells were labeled with anti-human CD3 (clone SK7)-APC (BD Bioscience) and NHP T cells were labeled with anti-human CD3 (clone SP34-2)-Pacific Blue (BD Pharmingen). SYTOX Blue (Invitrogen) or SYTOX Red (Invitrogen) were used to determine cell viability in human or NHP cells, respectively as follows: SYTOX Red/Blue was diluted to a final concentration of 5 nM in stain buffer. Cells were resuspended in 150 μL SYTOX Red/Blue and incubated for 15 minutes at room temperature protected from light. Samples were then processed by flow cytometry without washing.

Flow cytometric analysis was performed for all samples using an Attune NxT flow cytometer (ThermoFisher Scientific). Single-stained compensation controls and fluorescence-minus-one (FMO) controls were included for all analyses.

### B7-H3 construct characterization and creation of stable PG13 producer cell line

To generate the canine B7-H3-CAR construct, a previously developed B7-H3-specific single-chain variable fragment (scFv) [34] derived from the MGA271 monoclonal antibody [35] was subcloned into a canine CAR backbone containing a short hinge, canine CD28 transmembrane and costimulatory domains, and a canine CD3ζ signaling domain [8]. The final CAR plasmid was verified by sequencing.

GALV-pseudotyped B7-H3 CAR retrovirus was produced from stably transduced PG13 cells. Stable transduction was achieved as follows: Retroviral particles were generated by transient transfection of 293T cells (ATCC, CRL-11268) with the CAR-encoding vector, PeQ-Pam plasmid encoding MoMLV gag-pol, and a plasmid encoding the VSV-G envelope. Supernatant containing VSV-G-pseudotyped retroviral particles was collected, centrifuged, and filtered. Viral supernatant was then used to transduce PG13 producer cells. Transduced PG13 cells were stained for CAR expression using a F(ab′)_2_ fragment–specific antibody (polyclonal, Jackson ImmunoResearch Labs) and flow cytometrically sorted to isolate the top 10% of CAR-expressing cells. Sorted PG13 producer cells were expanded and used to generate GALV-pseudotyped retroviral supernatant for transduction of canine peripheral blood mononuclear cells (PBMCs).

### Creation of stable FLYRD18 CAR retrovirus producer cell line

RD114-pseudotyped B7-H3 CAR retrovirus was produced from stably transduced FLYRD18 cells. Stable transduction was achieved as follows: 8×10^5^ FLYRD18 cells (generously gifted by Dr. Gianpietro Dotti, UNC) were plated in each well of a 12-well tissue culture-treated well plate (Genesee) and cultured overnight in IMDM media (Life Technologies) supplemented with 10% FBS, 2 mmol/L GlutaMAX, and 100 U/mL Penicillin and 100 μg/mL Streptomycin (complete IMDM). The next day, cells were transfected with 1.25, 2.5, or 5 μg of purified B7H3-CAR plasmid DNA (pDNA) (prepared using ZymoPure midiprep kit according to manufacturer’s protocol) using Genejuice Transfection Reagent (Sigma) at varying doses. For each DNA dose, 1.5, 3, 6, and 9 μL of GeneJuice per μg of pDNA were evaluated, resulting in 12 distinct transfection conditions.

For each transfection, the respective amount of GeneJuice was added to 50 μL of non-supplemented IMDM media, gently mixed, and incubated at room temperature for 5 minutes. Then the GeneJuice and media mixture was dropwise added to the corresponding amount of pDNA and incubated at room temperature for 15 minutes. After incubation, the transfection solutions were each added to a respective FLYRD18-containing well of the 12 well plate. After gentle mixing, the 12-well plate was then placed back in the incubator.

After 48 hours, the media was collected and virus production was evaluated by transducing Jurkat cells in a similar manner to canine PBMCs with RetroNectin. The top 8 producing cells were maintained in culture in order to assess possible stable integration of the pDNA over time in the transfected cells. After approximately 2 weeks, CAR expression was evaluated by flow cytometry using αG4S linker antibody as a determinant of stable transduction. The top 3 cell lines (3.8-4.5% CAR positive cells) were maintained in culture and virus production was again measured after several days. The group producing the most virus was chosen for cell selection by flow cytometry. CAR^+^ FLYRD18 cells were single-cell sorted into four tissue culture treated 96 well plates (Genesee) and expanded. Wells with no growth, poor growth, or containing more than one colony were excluded and 48 remaining populations were selected for transfer to 24 well plates for expansion. After 3 days, virus production was evaluated and the top four virus producing cell lines were selected for cryopreservation.

### Virus Production

Retroviral particles were produced using PG13 or FLYRD18 retroviral producer cell lines maintained in DMEM media (Life Technologies) supplemented with 10% FBS, 2 mmol/L GlutaMAX, and 100 U/mL Penicillin and 100 μg/mL Streptomycin (complete DMEM). Stably transduced FLYRD18 cell line producing GFP gammaretrovirus was generously gifted by Gianpietro Dotti, UNC. Following three passages, culture medium was replaced with complete IMDM during the final passage. Viral supernatant from FLYRD18 cells was collected 20–24 hours after cultures reached 90-100% confluency. For PG13 cells, sodium butyrate (5 mM final concentration) was added 20–24 hours before viral collection to enhance viral production. Supernatants from both producer cell lines were collected, passed through a 0.45 μm vacuum filter, snap-frozen using a dry ice–ethanol bath, and stored at −80°C until use.

Virus titers were determined by transducing 10^6^ activated canine T cell enriched PBMCs as above. Two-fold serial dilutions ranging from 1X (undiluted) to 512X were used for transduction. Cells were expanded for 7 days and transduction efficiency was evaluated by flow cytometry. We selected cells with 5-20% transduction efficiency and calculated functional titer (transducing units (TU)/mL) using the following equation.

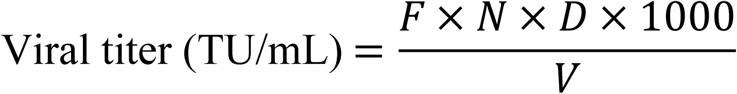

Where:

F = Fraction of transduced cells (e.g., 0.20 for 20%)
N = Number of target cells
D = Viral dilution factor
V = Volume of viral suspension added (µL)
1,000 = Conversion factor from µL to mL

Viral titers for each batch were as follows: GALV B7-H3 CAR retrovirus: 1.5 x 10^7^ TU/mL; RD114 B7-H3 CAR retrovirus: 4 x 10^5^ TU/mL; RD114 GFP retrovirus: 7 x 10^6^ TU/mL.

### Ethics Statement

Peripheral whole blood samples from dogs were obtained by the Laboratory Animal Resources veterinary team at North Carolina State University. All animal procedures were reviewed and approved by the Institutional Animal Care and Use Committee (IACUC) at the North Carolina State University College of Veterinary Medicine under Protocol Nos. 23-236 or 26-266. Blood collection and animal handling were conducted in accordance with institutionally approved animal care and use guidelines. Measures were taken throughout sample collection to limit stress and discomfort, including the use of appropriate handling and restraint practices. In particularly fearful animals, sedation with Gabapentin and Trazodone was utilized to alleviate anxiety. Donor dogs did not experience excessive or prolonged pain or distress that necessitated the administration of immobilizing sedation or anesthesia.

Peripheral whole blood samples from pigtail macaques (*Macaca nemestrina*) were obtained through the Tissue Distribution Program of the Washington National Primate Research Center (WaNPRC; now the Washington National Biomedical Research Center), University of Washington, Seattle, WA. Samples were collected by WaNPRC veterinary staff from animals designated for the Tissue Distribution Program, and no animals were used or procedures performed solely for the purposes of this study. All procedures were reviewed and approved by the University of Washington Institutional Animal Care and Use Committee (IACUC) under Protocol No. 4277-01. Animals were housed and cared for in accordance with the Guide for the Care and Use of Laboratory Animals, the Animal Welfare Act and Regulations, and the Public Health Service Policy on Humane Care and Use of Laboratory Animals at an AAALAC International-accredited facility (PHS Animal Welfare Assurance No. D16-00292).

Human peripheral whole blood samples were purchased from Gulf Coast Regional Blood Center (Houston, TX, USA) and obtained from anonymous, voluntary donors.

### Statistical analysis and figures

Flow cytometry data were analyzed using FCS Express 7 Research Edition software. Statistical analyses were performed using GraphPad Prism 9. All figures were generated using both FCS Express 7 Research Edition and GraphPad Prism 9.

## Results

### IL-2 promotes robust T cell expansion with less differentiated memory subsets

CAR T cell viability, expansion, and persistence are critical determinants of successful immunotherapy. Accordingly, we prioritized cytokine supplementation as an initial focus for improving these parameters. In humans, IL-2 drives robust T cell proliferation but generally yields terminally differentiated memory subsets [36], whereas IL-7 and IL-15 preserve high proportions of T_SCM_ and T_CM_ cells [29,36–38], which have been associated with better patient outcomes [21,22]. Whether these relationships hold in dogs is unclear, as only cursory studies have examined these cytokines in canine T cells [19,28].

To investigate the effects of cytokine supplementation on canine T cell activation and expansion, we evaluated varying concentrations of rhIL-2 compared to rhIL-7 and rhIL-15 during a stimulation period with plate-bound antibodies. In a concurrent experiment, cells activated in the same fashion were removed from stimulation after 3 days and expanded for 4 days in their respective cytokines/concentrations. Memory phenotypes were assessed at day 3 and 5 of the stimulation period, and after 4 days of expansion post-stimulation. Media containing no cytokines was used as a negative control.

During the stimulation period, there was no observable proliferation difference between any of the tested cytokines or cytokine concentrations and total expansion was comparable to the negative control group (Fig 1A). We also noted that during stimulation, exogenous cytokines had little impact on memory subset differentiation (Fig 1B and S1 Fig). Taken together, these results suggest that endogenous cytokine release in stimulated T cells may be the major contributor to proliferation and phenotype determination.

**Fig 1.**
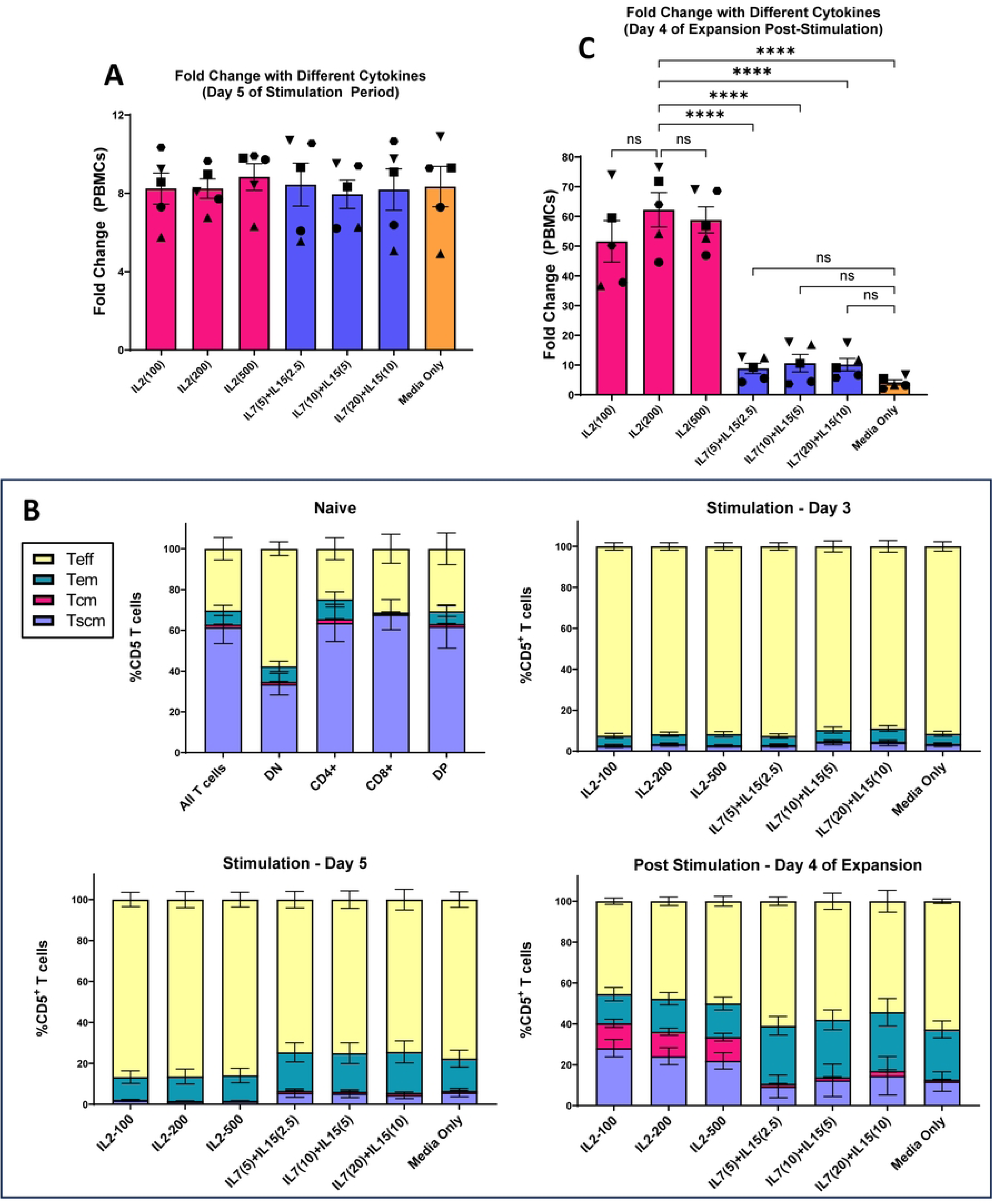
IL-2 Improves expansion of canine T cells after stimulation while maintaining a less differentiated memory phenotype. PBMCs from five canine donors were cultured with different concentrations of rhIL-2, rhIL-7 and rhIL-15, or without cytokines (media only). A) Expansion was evaluated after a 5-day stimulation period. B) Memory phenotypes in CD5^+^ T cells were also evaluated prior to stimulation (naive; DN=CD4^-^CD8^-^ double negative, DP=CD4^+^CD8^+^ double positive), after a 3- and 5-day stimulation period, and after 4 days of expansion post-3-day stimulation period. C) Fold-expansion after removal from stimulation was evaluated on day 4. Horizontal lines indicate mean values, and error bars represent standard error of the mean. Pairwise statistical analysis was performed by one-way ANOVA with Tukey’s multiple means comparison; * p<0.05, ** p<0.01, *** p<0.001, **** p<0.0001, ns not significant. Different symbols represent individual donors.

After cells were removed from stimulation, however, the cytokine treatments induced dramatic differences between groups. As expected from previous publications [20,26], IL-2 supplementation resulted in robust expansion with the highest fold-increase seen at 200 IU/mL. All doses resulted in significantly higher expansion than any of the IL-7+IL-15 doses. While IL-7+IL-15 increased impact on expansion compared to the negative control, it failed to reach statistical significance (Fig 1C). During expansion, the tested cytokines also played a crucial role in T cell memory differentiation (Fig 1B). Not only did IL-2 supplementation result in approximately double the proportion of T_SCM_ cells compared to IL-7+IL-15, but the combined T_SCM_ and T_CM_ cells accounted for as much as 40% of all T cells. This was much higher than the peak of 17% seen with IL-7+IL-15, which was not distinguishable from the negative control.

IL-2 also contributed to maintenance of a morphologically larger T cell population, consistent with an activated phenotype. IL-7+IL-15 improved maintenance of this population more than the negative control, but to a far lesser degree than IL-2 (Fig 2A). IL-7+IL-15 significantly improved T cell viability compared to the negative control at all doses. However, IL-2 supplementation resulted in significantly improved viability over IL-7+IL-15 for all but the highest dose (Fig 2B-D).

**Fig 2.**
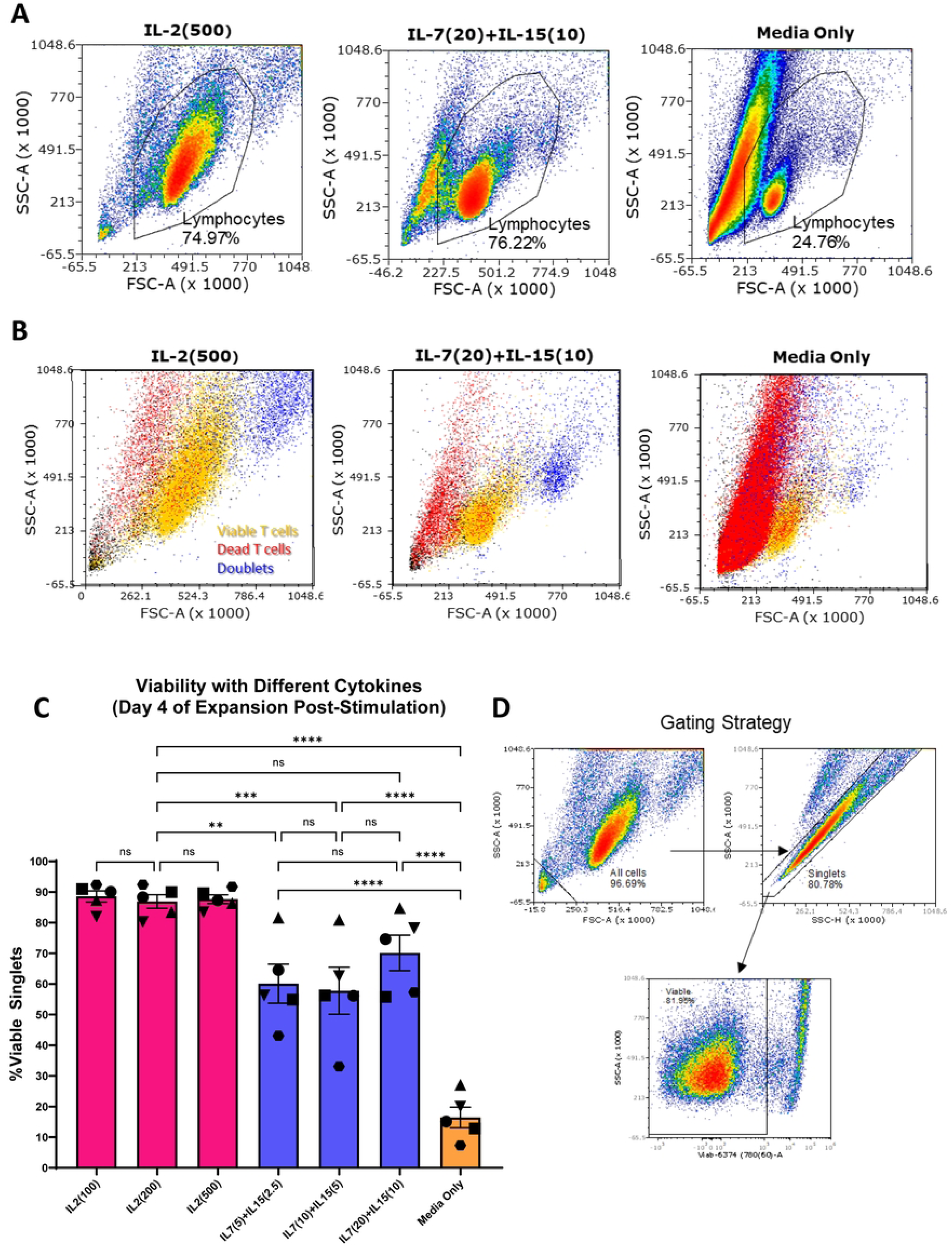
IL-2 improves canine T cell viability while maintaining an activated morphology. PBMCs from five canine donors were activated with stimulatory antibodies for 3 days in the presence of different concentrations of rhIL-2, rhIL-7 and rhIL-15, or without cytokines (media only). Cells were then expanded post-stimulation for 4 days under the same cytokine conditions. A) On day 4, flow cytometry was used to evaluate changes in cell size as an indicator of an activated T cell phenotype. In these representative plots, lymphocytes were gated in SSC/FSC plots and leftward shift on FSC axis indicated a decrease in cell size consistent with a loss in activated phenotype. B) Backgating was used to highlight differences in the experimental groups, where orange indicates CD5^+^ T cells, blue indicates doublets, and red indicates non-viable cells. C) Viability was quantified using flow cytometry where viable singlets were compared across groups on day 4. Horizontal lines indicate mean values, and error bars represent standard error of the mean. Pairwise statistical analysis was performed by one-way ANOVA with Tukey’s multiple means comparison; ** p<0.01, *** p<0.001, **** p<0.0001, ns not significant. Different symbols represent individual donors. D) Representative plot for gating strategy used to assess viability.

Taken together, supplementation with IL-2 after antibody stimulation resulted in improved viability, increased T cell proliferation and a more favorable memory profile while maintaining an activated state when compared to IL-7+IL-15. IL-7+IL15 improved viability but did not meaningfully affect memory phenotype or expansion compared to cytokine free conditions. For subsequent experiments, IL-2 dosing was standardized at 200 IU/mL due to its slight expansion improvement compared to other IL-2 doses.

### Two-day stimulation period with high cell density yields higher transduction efficiency

Having identified cytokine culture conditions that supported favorable canine T cell expansion and differentiation, we next evaluated strategies to improve transduction efficiency. Activated T cells actively undergoing mitosis are more susceptible to retroviral transduction due to breakdown of the nuclear envelope [39] and activation occurs early in dogs after stimulation with plate-bound antibodies (S2 Fig). While we established optimal stimulation methods in our previous work [20], we did not establish how the duration of stimulation affects transduction success. To determine this effect, PBMCs from four healthy canine donors were stimulated with plate-bound antibodies for 2–5 days before transduction. For each group 10^6^ PBMCs were transduced and allowed to expand for 7 days. Transduction efficiency was assessed at 2, 4, and 7 days post-transduction, and cellular expansion was evaluated on day 7. A non-transduced control group was included for comparison.

For cells transduced after 3 or more days of stimulation, transduction efficiency stayed consistent from day 2 of expansion onwards. However, for cells stimulated for only 2 days, transduction increased during expansion, ultimately exceeding transduction of all other groups (Fig 3A). The stimulation period had a direct impact on expansion potential (Fig 3B).

**Fig 3.**
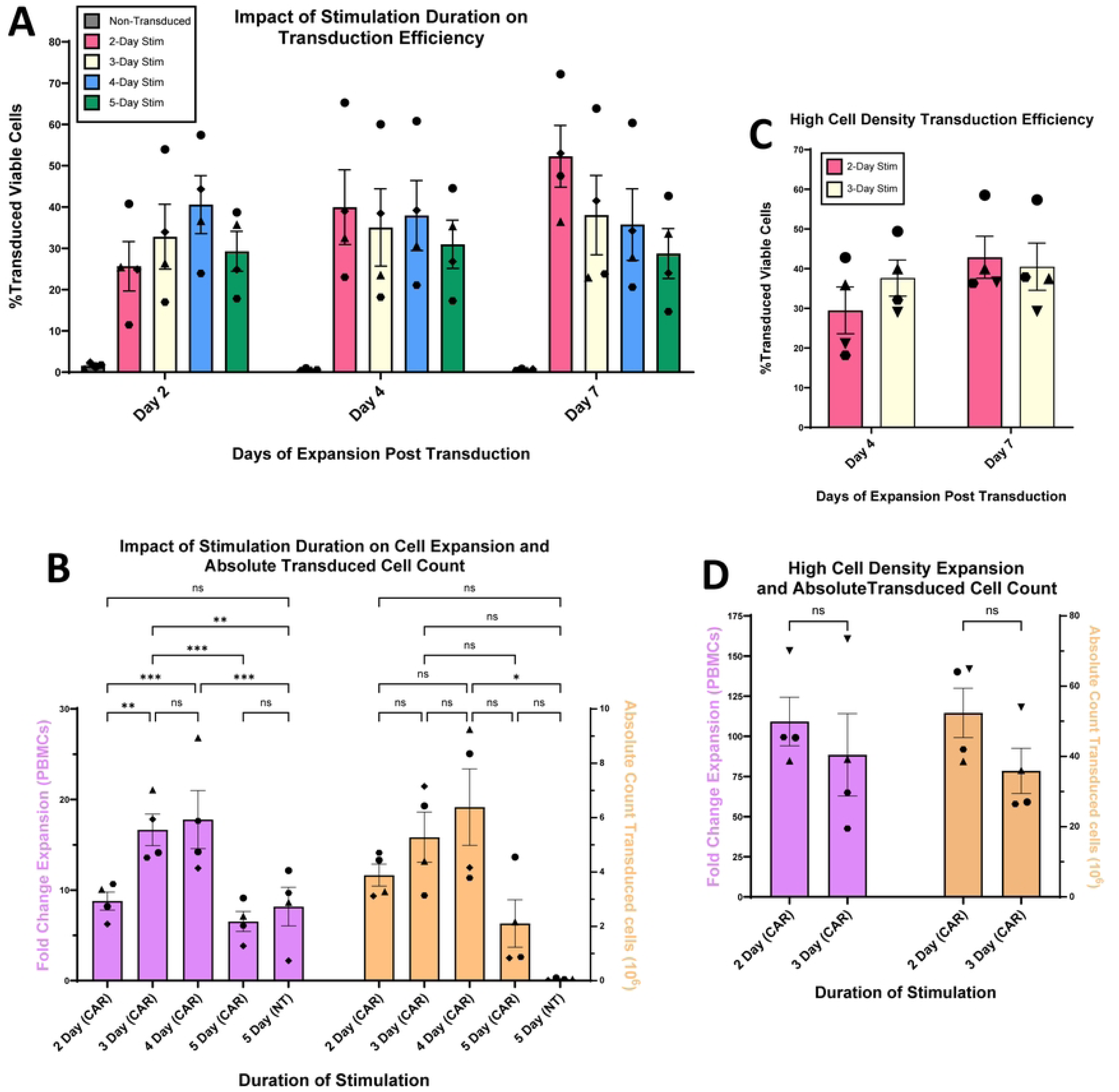
At higher cell densities, stimulation for 2 days improves transduction and expansion rates in canine T cells. A) PBMCs from four canine donors were activated with stimulatory antibodies for 2-5 days. After removal from stimulation, T cell-enriched PBMCs were transduced and transduction rate was evaluated by flow cytometry for viable lymphocytes on days 2, 4, and 7. Non-transduced cells (NT) were stimulated for 5 days, but were not transduced. B) On day 7 after transduction, expansion was evaluated for all groups (purple bars) and the absolute number of transduced cells was calculated (orange bars). C) PBMCs from four canine donors were again activated with stimulatory antibodies for 2 or 3 days. Initial PBMC cell seeding density was increased. Transduction efficiency was evaluated by flow cytometry for viable lymphocytes on days 4 and 7. D) On day 7 after transduction, expansion of the two groups was evaluated (purple bars) and the absolute number of transduced cells was calculated (orange bars). For all figures, horizontal lines indicate mean values, and error bars represent standard error of the mean. Statistical analysis was performed by two-way ANOVA with Tukey’s multiple means comparison; * p<0.05, ** p<0.01, *** p<0.001, **** p<0.0001, ns not significant. Different symbols represent individual donors.

Stimulating for 2 days was insufficient for robust cell expansion, resulting in reduced cell numbers. Poor expansion was also seen in cells stimulated for 5 days, whether transduced or not, suggesting that the 5-day stimulation itself is impairing proliferation. Taking both transduction efficiency and cell expansion together, the 3-day and 4-day stimulation resulted in significantly improved expansion and thus a higher absolute number of transduced cells (Fig 3B).

Phenotypic analysis revealed that the 2-day stimulation group had a significantly higher proportion of CD4^+^ T cells than all other groups (S3 Figs A and B and S4 Fig). This finding is particularly relevant given the growing recognition of CD4^+^ CAR T cells as important contributors to antitumor activity and long-term therapeutic responses [22]. In addition, 2-and 3-day stimulation groups had significantly higher proportions of T_SCM_ cells, with a sharp decline observed in 4- and 5-day stimulation groups (S3 Figs C and D).

With longer stimulation periods negatively impacting T cell memory subsets, we focused on boosting expansion in cells stimulated for 2 or 3 days. Increasing cell density has been associated with better proliferation in human T cells [40]. Thus we tested whether higher PBMC density during 2 or 3 days of stimulation would promote improved T cell expansion and CAR T cell production. As above, after stimulation, 10^6^ cells from each group were transduced and allowed to expand for 7 days. Increasing PBMC seeding density during the stimulation period not only dramatically increased expansion rates for both groups compared to the previous experiment, but also abrogated the poor expansion previously seen with 2 days of stimulation. After 7 days of expansion, 2-day stimulation again produced the best transduction efficiency (Fig 3C), but this time it also resulted in superior expansion translating to higher absolute numbers of transduced cells (Fig 3D).

With higher transduction, improved expansion, and a high T_SCM_ proportionality, we deemed a 2-day stimulation period with high PBMC seeding density to be ideal activation conditions for canine T cell transduction. In light of these findings, this protocol was used for all subsequent transduction experiments.

### Phosphate deprivation does not enhance GALV-mediated transduction

Despite our process improvements, canine T cell transduction with GALV-pseudotyped CAR retrovirus consistently plateaued near 50%. We therefore sought methods to further enhance transduction to maximize our CAR T cell yield. Phosphate deprivation has been reported to increase GALV retroviral transduction efficiency in human and non-human primate (NHP) lymphocytes through triggering the upregulation of the phosphate transporter protein PiT1 (SLC20a1), the cellular entry receptor for GALV [41].

We evaluated whether phosphate deprivation could improve canine T cell transduction. PBMCs from four healthy donors were stimulated according to our improved protocol. 16 hours or 6 hours prior to transduction, media was replaced with phosphate free media containing 3% FBS. A control group was maintained in standard media prior to transduction (S5 Fig A). Cells were transduced and expanded in standard media. In this instance, phosphate deprivation did not improve transduction (S5 Fig B).

Because FBS contains inorganic phosphate, a similar experiment was performed using phosphate-free medium without serum to further restrict phosphate availability. At transduction, cells were either returned to standard media or underwent continued phosphate/serum deprivation overnight (Fig 4A). All groups were expanded the following day in standard media. Again, phosphate/serum deprivation prior or during transduction did not enhance transduction efficiency in canine T cells (Fig 4B) and ultimately failed to improve the absolute number of transduced cells (Fig 4C).

**Fig 4.**
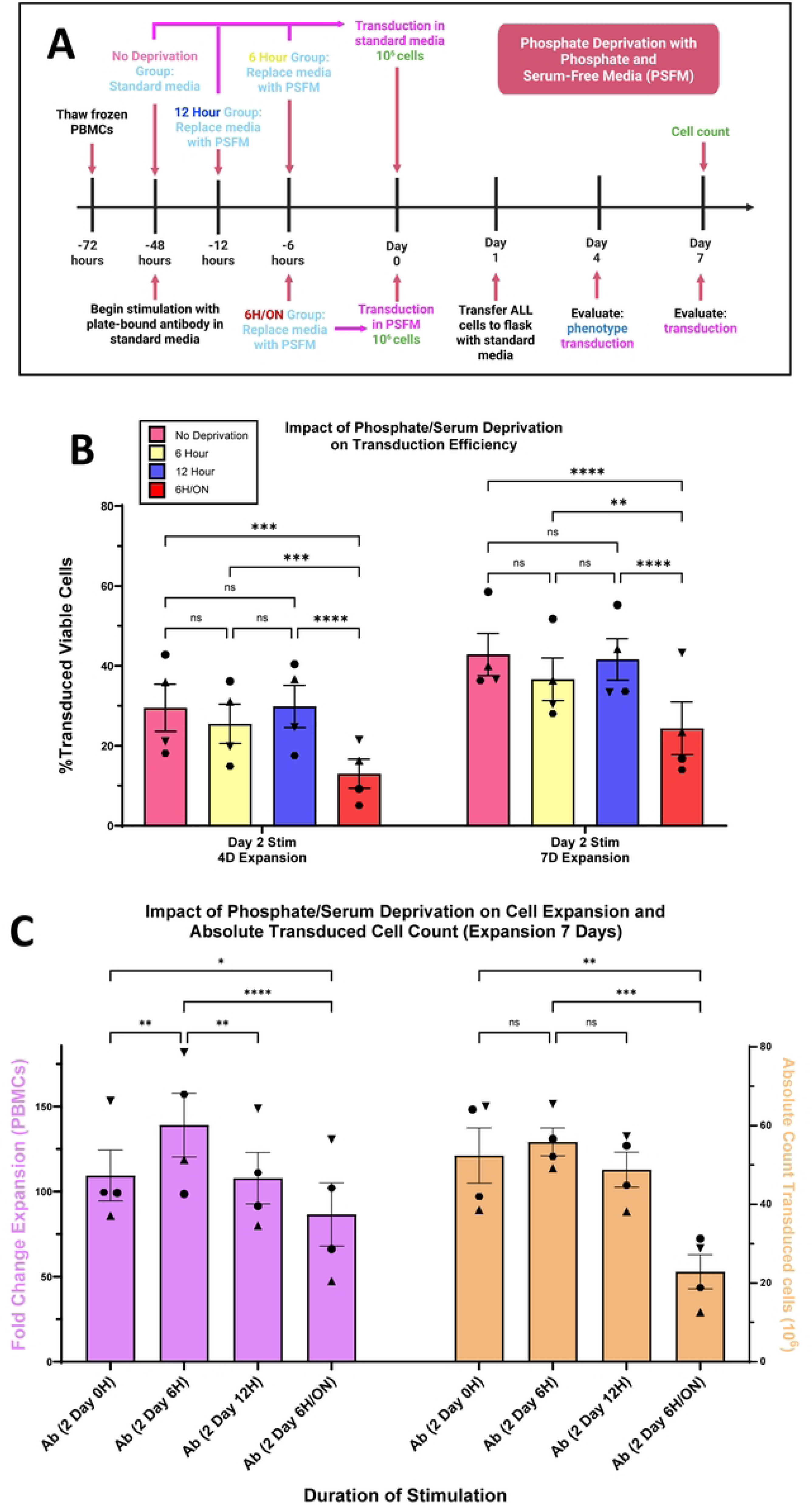
During the transduction period, serum/phosphate deprivation diminishes transduction efficiency with GALV-pseudotyped retrovirus, while deprivation prior to transduction has no impact. A) Schematic of the experiment design. PBMCs from four canine donors were activated with stimulatory antibodies for 2 days. On the day of transduction, standard 10% FBS T cell media was replaced with serum and phosphate free media 12 and 6 hours before transduction. The “No Deprivation” group was maintained in standard media. All groups were returned to standard media during the transduction period, except for the “6H/ON” group. For this group, standard media was replaced with serum/phosphate free media for 6 hours before transduction and cells were maintained in this media overnight during the transduction period. The day after transduction, all cells were then expanded in IL-2 enriched standard 10% FBS media and B) transduction was evaluated by flow cytometry in viable cells on days 4 and 7. C) Expansion after transduction was evaluated for all groups (purple bars) and the absolute number of transduced cells was calculated (orange bars). For all figures, horizontal lines indicate mean values, and error bars represent standard error of the mean. Statistical analysis was performed by two-way ANOVA with Tukey’s multiple means comparison; * p<0.05, ** p<0.01, *** p<0.001, **** p<0.0001, ns not significant. Different symbols represent individual donors.

In the group deprived of phosphate/serum overnight during the transduction period, we observed a significant reduction in expansion and, more notably, in transduction efficiency. This suggests that either phosphate or components of FBS are important for virus particle/T cell interactions during the transduction process.

### FBS and cholesterol improve canine T cell transduction with GALV and RD114 retrovirus

To further investigate the influence of serum on canine T cell transduction, activated PBMCs from four healthy donors were transduced with GALV retrovirus in media containing increasing concentrations of FBS (0–50%). Transduction in serum free media (SFM) was significantly decreased compared to media with serum (Fig 5A). Transduction increased with increasing FBS concentration up to 30% FBS with no added benefit above 30% (Fig 5A). As previous studies have demonstrated a correlation between the number of integrated viral vector copies per cell (vector copy number, VCN) and protein expression measured by fluorescence intensity [42–44], we also evaluated median fluorescence intensity (MFI) at each FBS concentration. Increasing FBS concentration did not alter MFI among transduced cells, indicating that FBS may facilitate transduction of otherwise difficult-to-transduce cells rather than increase the number of viral copies integrated into individual cells (Fig 5B).

**Fig 5.**
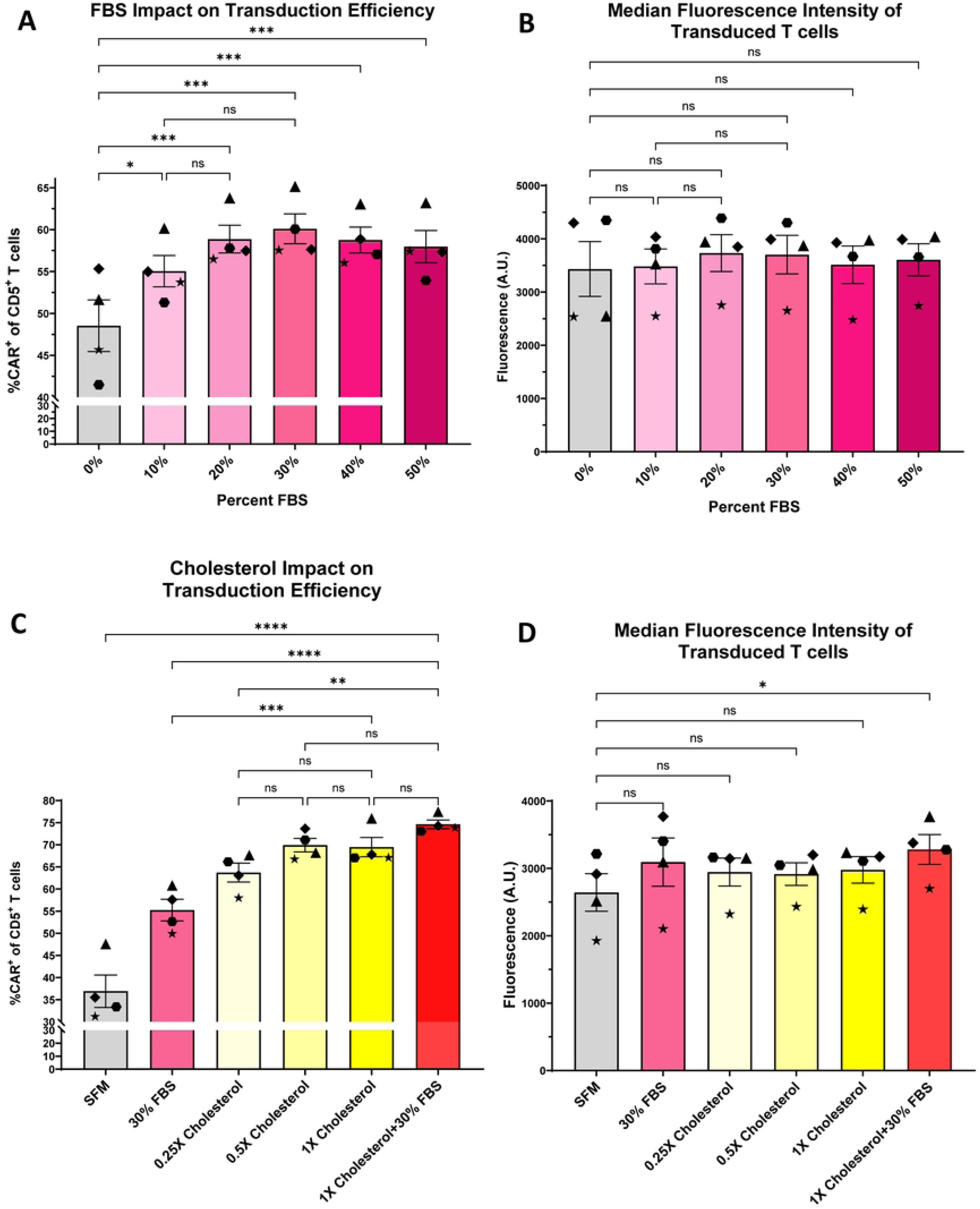
High concentration FBS and cholesterol enhance canine T cell transduction with GALV-pseudotyped retrovirus. A) PBMCs from four canine donors were activated with stimulatory antibodies, then transduced with GALV-pseudotyped CAR retrovirus. During the transduction period, standard 10% FBS T cell media was replaced with media containing different concentrations of FBS and cells were cultured overnight. The following day, cells were transferred to standard 10% FBS media and expanded. Transduction was evaluated on day 7. B) The intensity of CAR signaling using median fluorescence intensity (MFI) in transduced CD5^+^ T cells was used as an indicator of virus particle/cell interaction. C) The experiment was repeated, but during the transduction period, standard media was replaced with media containing 0% FBS (SFM), 30% FBS with or without cholesterol, and different concentrations of cholesterol. Cells were maintained overnight before being transferred to standard media the following day for expansion. Transduction and D) MFI were evaluated on day 7. For all figures, horizontal lines indicate mean values, and error bars represent standard error of the mean. Pairwise statistical analysis was performed by one-way ANOVA with Tukey’s multiple means comparison; * p<0.05, ** p<0.01, *** p<0.001, **** p<0.0001, ns not significant. Different symbols represent individual donors.

As cholesterol in the cell membrane has been shown to be necessary for viral particle endocytosis/endosomal escape [45] and is a major component of FBS, we next investigated cholesterol as a primary driver of serum’s transduction enhancement. Activated PBMCs were transduced in media containing increasing concentrations of cholesterol (0.25X, 0.5X, and 1X based on Gibco 250X Cholesterol). Additional groups included serum-free media (SFM), 30% FBS, and 1X cholesterol combined with 30% FBS. Cholesterol at all concentrations significantly improved transduction over SFM and 30% FBS, with a nearly 2-fold peak enhancement over SFM. Interestingly, 1X cholesterol+30% FBS improved transduction over cholesterol alone and reduced donor variability (Fig 5C). When MFI was evaluated, this group alone had significantly higher signal than SFM transduction, raising the possibility that this combination increases interactions between viral particles and T cells, potentially resulting in greater viral integration and CAR expression per cell. (Fig 5D). These transduction enhancements were verified with a repeat experiment (S6 Fig). Taken together, these findings demonstrate that cholesterol supplementation to media at the time of transduction can dramatically boost GALV-mediated retroviral transduction efficiency. The further enhanced transduction in combination with high concentration FBS implies the effects are additive, indicating there are other components in FBS that contribute to enhanced transduction beyond cholesterol/lipid alone.

We further investigated these effects across a range of viral concentrations using serial dilutions of GALV-pseudotyped CAR retrovirus. Two-fold serial dilutions ranging from 1X (undiluted; MOI=15) to 128X were used to transduce activated T cells cultured in either 1X cholesterol+30% FBS or 10% FBS (our standard transduction protocol). Once again, 1X cholesterol+30% FBS substantially increased transduction over standard 10% FBS transduction (77% versus 46%, respectively in undiluted virus) and significantly improved transduction across all dilutions. Remarkably, 16-fold less virus was needed with cholesterol/FBS transduction to achieve the same efficiency as standard FBS transduction using undiluted virus (Fig 6).

**Fig 6.**
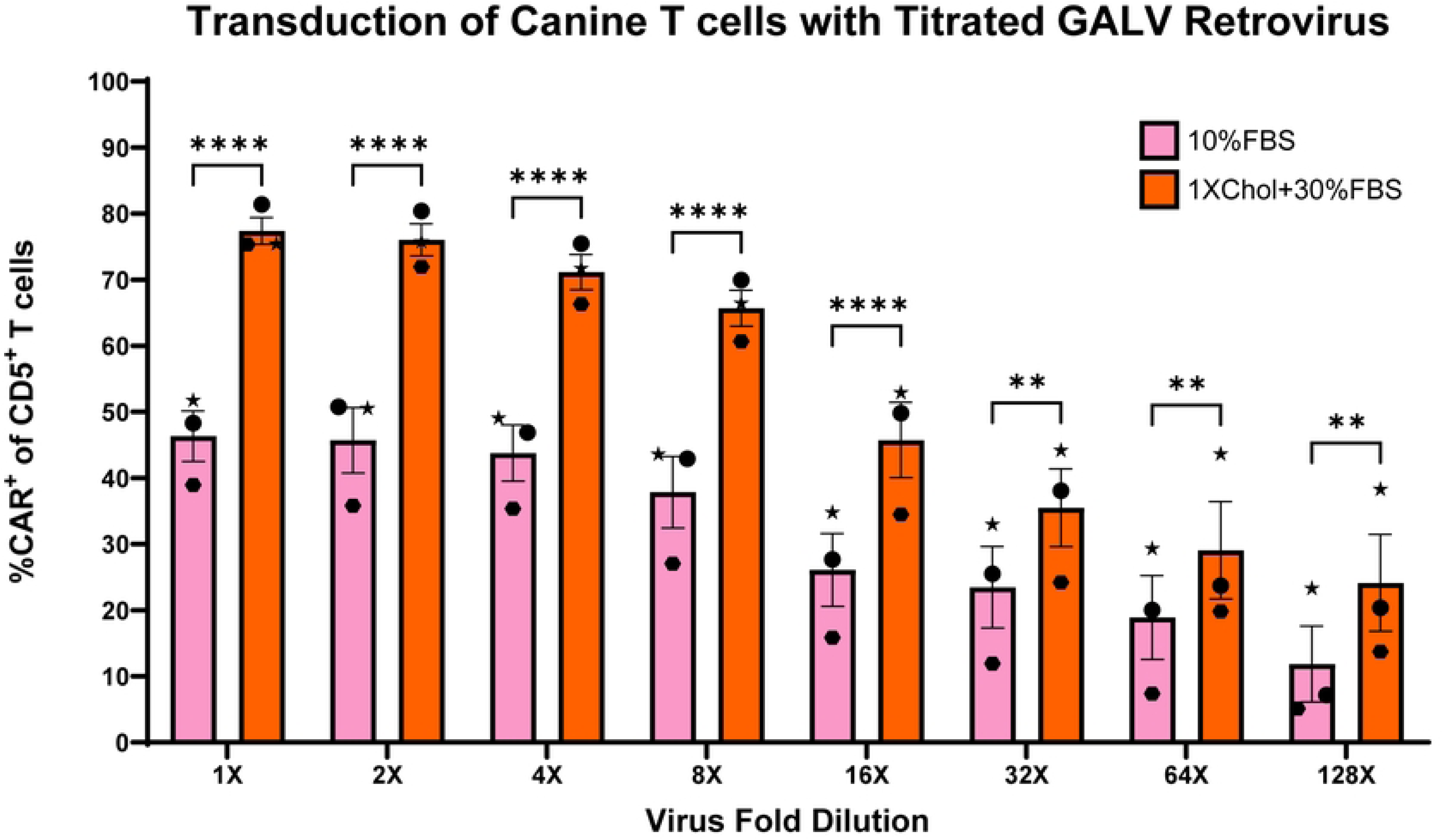
16-fold less virus with 1X Cholesterol+30% FBS is required to achieve the same transduction as standard media. After activation, PBMCs from three canine donors were transduced with GALV pseudotyped CAR retrovirus in dilutions ranging from 1X (undiluted; MOI=15) to 128X. Transduction was evaluated on day 7 in CD5^+^ T cells. Statistical significance between culture conditions at each virus dilution was assessed by two-way repeated-measures ANOVA followed by Šídák’s multiple-comparisons test. **P < 0.01, ****P < 0.0001. Different symbols represent individual donors.

To determine if this effect could be reproduced with other viral envelopes, activated T cells were transduced as previous in media with 10% FBS, 1X cholesterol, or 1X cholesterol+30% FBS with low-titer RD114-pseudotyped CAR retrovirus (MOI=0.4). We noted a nearly 2-fold increase in transduction in the presence of cholesterol, which was further enhanced in combination with 30%FBS (Fig 7A).

**Fig 7.**
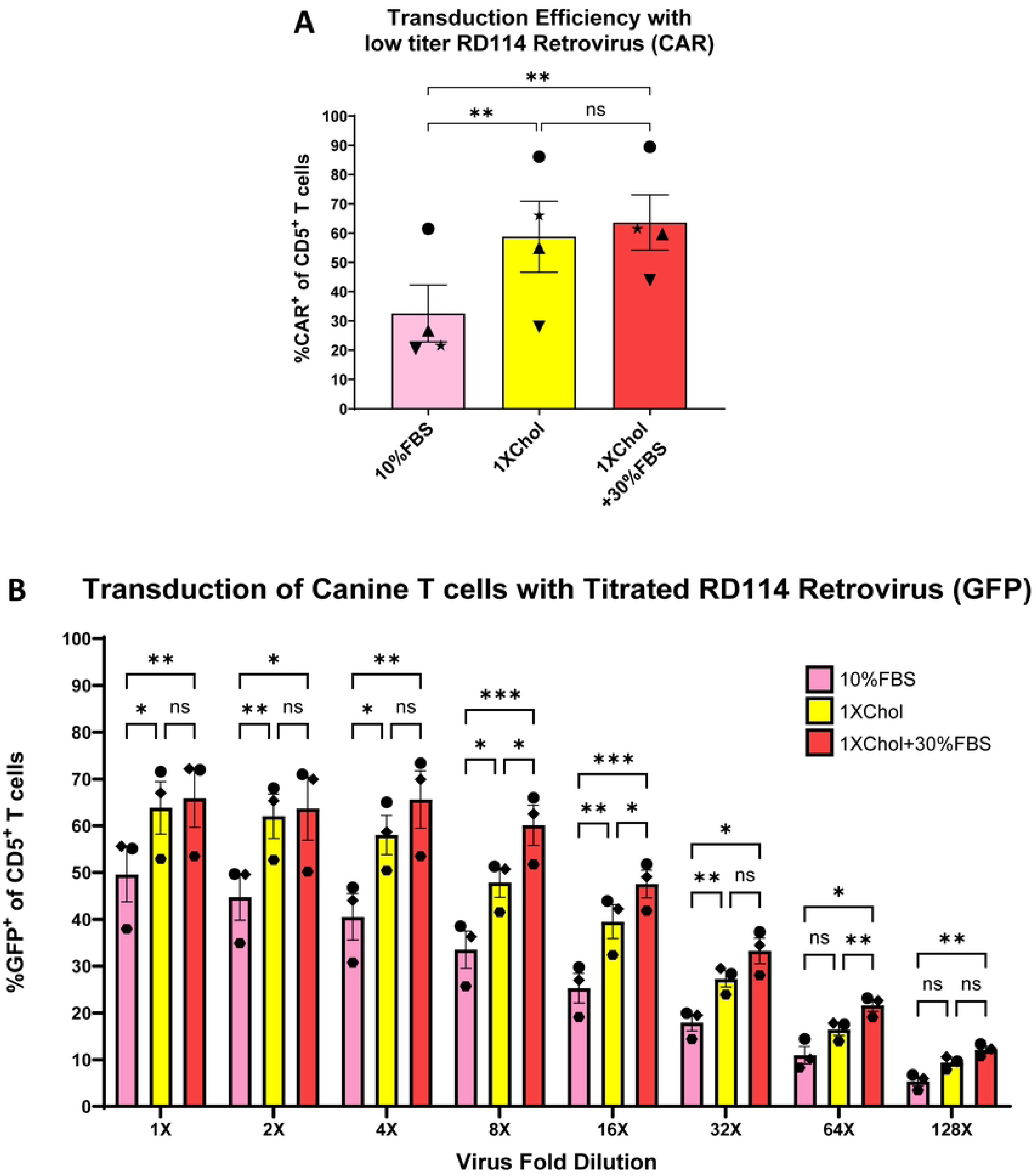
High concentration FBS with cholesterol enhances canine T cell transduction using RD114-pseudotyped retrovirus. A) PBMCs from four canine donors were activated with stimulatory antibodies, then transduced using a low-titer RD114-pseudotyped CAR retrovirus (MOI=0.4). During the transduction period, cells were either maintained in standard 10% FBS T cell media or transferred to media containing 1X cholesterol with or without 30% FBS. The following day, cells were transferred to 10% FBS media, expanded, and transduction efficiency was evaluated on day 7 in CD5^+^ T cells. B) Using the same culture conditions, PBMCs from three canine donors were transduced with RD114-pseudotyped GFP retrovirus in dilutions ranging from 1X (undiluted; MOI=7) to 128X. Transduction efficiency was evaluated in CD5^+^ T cells after 7 days. Statistical significance between culture conditions at each virus dilution was assessed by two-way repeated-measures ANOVA followed by Šídák’s multiple-comparisons test. *P < 0.05, **P < 0.01, ***P < 0.001; ns, not significant. For A) and B), horizontal lines indicate mean values, and error bars represent standard error of the mean. Different symbols represent individual donors.

We then transduced activated T cells using serial dilutions of high-titer RD114-pseudotyped GFP retrovirus under the same culture conditions (1X; MOI=7). Again, cholesterol with and without high concentration FBS significantly improved transduction, with the combination enhancing the effect. This effect was most apparent at 4X dilution and beyond where there is a clear divergence in transduction efficiency using high concentration FBS with cholesterol compared to cholesterol alone. As seen with GALV, 16-fold less RD114 virus was needed with cholesterol/FBS to achieve the same transduction in 10% FBS with undiluted virus. With cholesterol alone, 8-fold less virus was needed (Fig 7B).

### High concentration FBS and Cholesterol improve canine T cell expansion potential

Given the apparent synergistic effects of high-concentration FBS and cholesterol on canine T cell transduction, we further investigated their effects on cellular expansion. As FBS and cholesterol appear to enhance transduction through distinct mechanisms, bovine serum albumin (BSA), the predominant protein component of serum, was included in the analysis to determine whether it was primarily responsible for the transduction-enhancing effects of FBS. Activated PBMCs were transduced in media containing 10% or 30% FBS, BSA, or 1X cholesterol, alone or in combination with 30% FBS or BSA. Transduction efficiency and cellular expansion were subsequently evaluated.

Again, cholesterol dramatically improved transduction, which was again enhanced by the addition of high concentration FBS. This combination resulted in significantly higher transduction rates than all but the cholesterol alone treatment group. In contrast, adding BSA to cholesterol paradoxically negated much of cholesterol’s enhancement (Fig 8A and S7 Fig). Regarding expansion, 1X cholesterol alone produced a similar expansion profile to 10% or 30% FBS alone. However, combining 1X cholesterol with 30% FBS produced a synergistic effect, significantly increasing expansion and nearly doubling cell counts compared with either component alone. Conversely, BSA, with or without cholesterol, resulted in the lowest expansion (Fig 8B and C).

**Fig 8.**
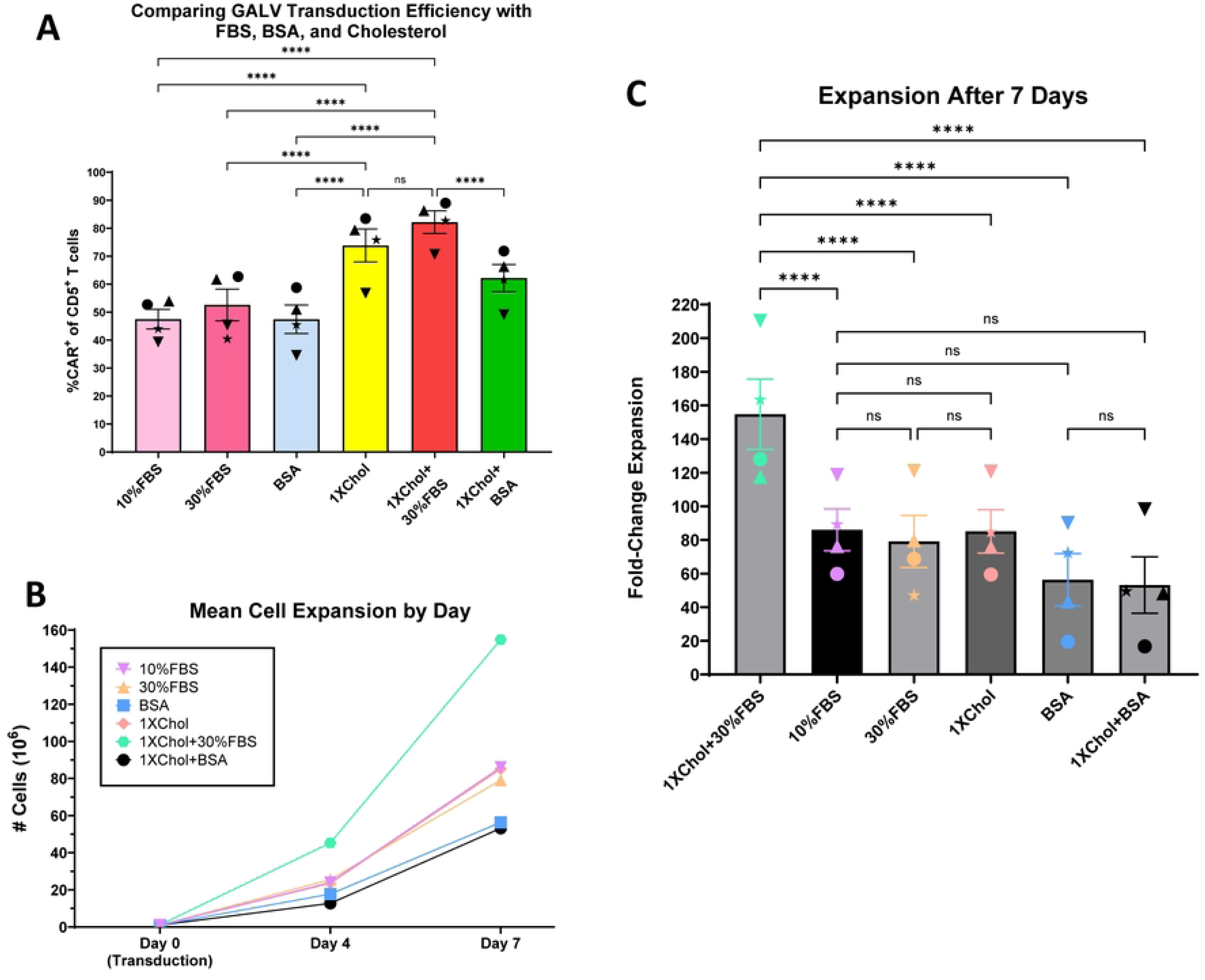
High concentration FBS and cholesterol work synergistically to enhance canine T cell transduction and expansion. PBMCs from four canine donors were activated with stimulatory antibodies, then 10^6^ cells were transduced with GALV-pseudotyped CAR retrovirus. During the transduction period, cells were either maintained in standard 10% FBS media or transferred to media containing 1X cholesterol, 30% FBS (+/- cholesterol), or BSA (+/- cholesterol). The next day, cells were transferred to standard media with IL-2 and expanded. A) Transduction was evaluated on day 7, B) cell counts were performed on days 4 and 7, and C) fold-change expansion was determined on day 7. For A) and C) horizontal lines indicate mean values, and error bars represent standard error of the mean. Pairwise statistical analysis was performed by one-way ANOVA with Tukey’s multiple means comparison; **** p<0.0001, ns not significant. Different symbols represent individual donors.

### FBS/cholesterol enhances T cell proliferation while preserving memory phenotype

With the marked enhancement of expansion seen with high concentration FBS and cholesterol, we were concerned that this may be driving terminal differentiation of memory T cell subsets, which would negatively affect persistence *in vivo* [24]. To explore this, we transduced and expanded T cells as previous using cholesterol (0.25X, 0.5X, and 1X) with high concentration FBS. Transduction with standard 10% FBS was used as a control. Cells were expanded in standard media, and after 4 days we assessed expansion rates and screened for any T cell phenotype changes; transduction was determined on day 7. High concentration FBS with all doses of cholesterol significantly enhanced proliferation compared to standard media, with highest expansion seen with 0.5X cholesterol, and lowest variability with 1X cholesterol (Fig 9A).

**Fig 9.**
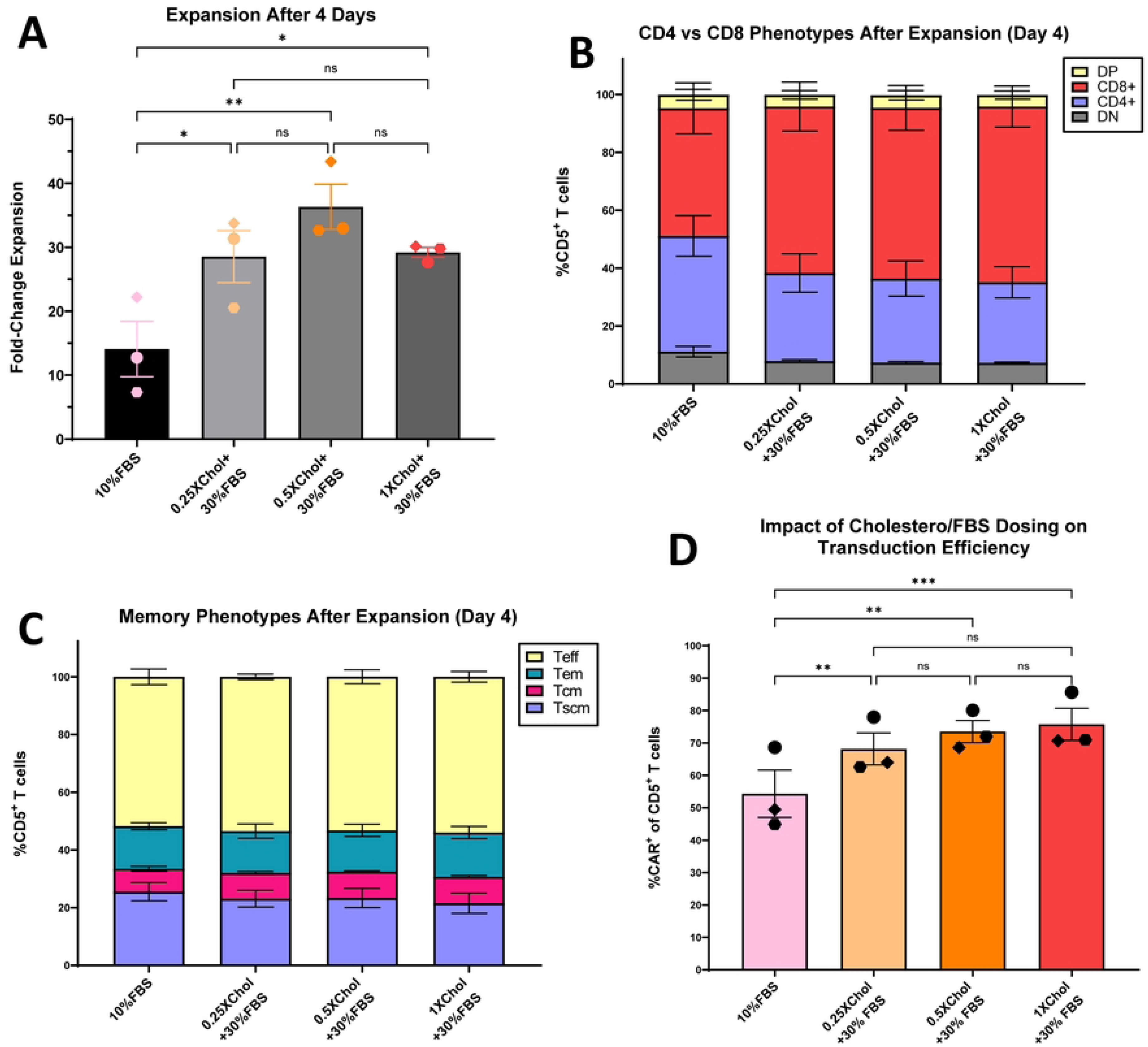
High concentration FBS with cholesterol preferentially expands CD8^+^ T cell subsets while maintaining a less differentiated memory phenotype. Activated PBMCs from three canine donors were transduced with GALV-pseudotyped CAR retrovirus. During the transduction period, cells were either maintained in standard 10% FBS T cell media or transduced in media containing 30% FBS with different concentrations of cholesterol. The following day, cells were transferred to standard 10% FBS for expansion. On day 4, A) fold-change expansion was determined as well as B) CD4 vs. CD8 subsets and C) memory subsets in the CD5^+^ T cell population. D) On day 7, transduction was compared across groups. For A) and D) pairwise statistical analysis was performed by one-way ANOVA with Tukey’s multiple means comparison; * p<0.05, ** p<0.01, *** p<0.001, ns not significant. Different symbols represent individual donors.

Across all cholesterol doses, high concentration FBS/cholesterol favored CD8^+^ T cell expansion, resulting in a higher CD8:CD4 ratio compared to standard media (Fig 9B). Robust expansion with FBS/cholesterol appeared to have minimal impact on T cell memory subsets, and a robust T_CM_ and T_SCM_ population was maintained across all cholesterol doses (Fig 9C). The proportions of T_SCM_ + T_CM_ cells or T_CM_ alone were not significantly different across groups (S8 Fig A and B). While a statistically significant reduction in T_SCM_ proportion was observed compared to the standard media group, the difference was only 4% (S8 Fig C). Across cholesterol doses, there was no significant increase in terminally differentiated T_EFF_ cells (S8 Fig D). All cholesterol doses resulted in higher transduction rates compared to standard media but were broadly similar to each other, with a slight increasing trend as the dose increased (Fig 9D).

Taken together, high concentration FBS with cholesterol during transduction enhanced transduction efficiency and subsequent expansion regardless of cholesterol dose. This enhanced expansion favors CD8^+^ T cell subsets without driving terminal differentiation of T cells.

### High concentration FBS and cholesterol prevent transduction impedance at high temperatures

Dogs maintain a higher basal body temperature than humans, averaging around 38.8°C. Thus, it has been reported that culturing canine T cells at 38.8°C enhances activation, transduction, and expansion compared to 37°C [20,32]. However, we found that temporarily reducing the temperature to 37°C during overnight transduction further improved transduction efficiency, potentially reflecting reduced viral stability and infectivity at higher temperatures. Andreadis and Palsson proposed that FBS may improve virus particle stability [46]. To explore this as one possible contributor to enhanced transduction, we performed transduction either at 37°C or 38.8°C and then expanded cells at 38.8°C.

Overnight transduction at 38.8°C reduced transduction efficiency across most culture conditions (Fig 10). Although 10% FBS, 30% FBS, and cholesterol supplementation exhibited comparable reductions, BSA alone and 1X cholesterol+30% FBS showed minimal temperature-associated declines. Notably, the cholesterol/FBS combination maintained the highest absolute transduction efficiency with reduced inter-donor variability (S9 Fig A). In contrast, combining cholesterol with BSA resulted in a significant temperature-associated reduction in transduction efficiency and the greatest relative decline among treatment groups (S9 Fig B). Together, these findings indicate that cholesterol and high-concentration FBS supplementation may mitigate temperature-associated reductions in retroviral transduction efficiency.

**Fig 10.**
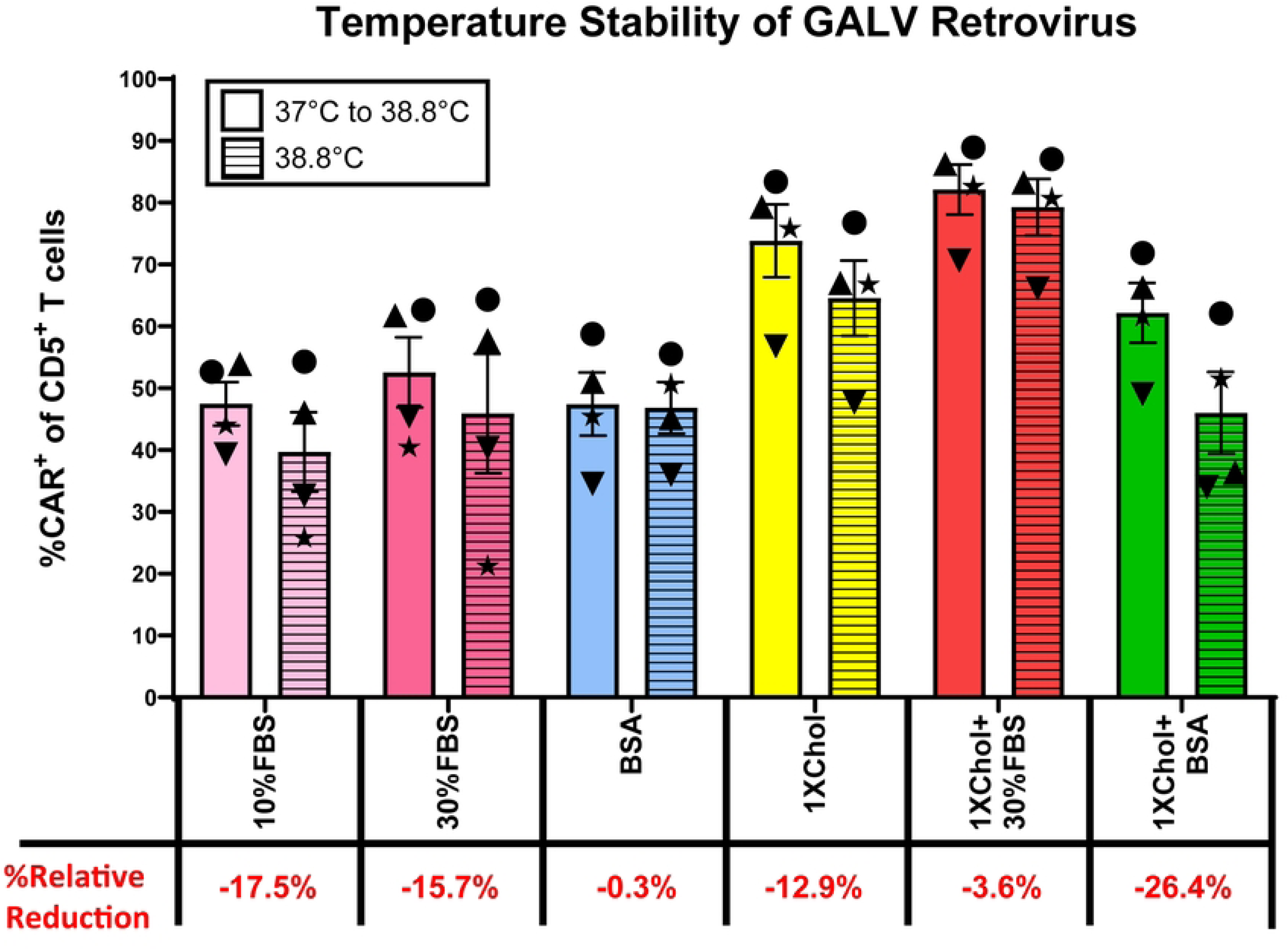
High concentration FBS and cholesterol work synergistically to prevent transduction impedance at high temperatures. PBMCs from four canine donors were activated with stimulatory antibodies, then transduced with GALV-pseudotyped CAR retrovirus. During the transduction period, cells were either maintained in standard 10% FBS T cell media or transferred to media containing 1X cholesterol, 30% FBS (+/-cholesterol), or BSA (+/-cholesterol). Cells were cultured at 37°C or 38.8°C overnight. The next day, cells were transferred to standard media with IL-2 and expanded at 38.8°C. Changes in transduction were evaluated under these different temperature culture conditions. Transduction rates at day 7 were compared between groups transduced at 37°C and 38.8°C. Relative reduction in mean transduction efficiency was compared across groups (red font).

### The FBS/cholesterol transduction enhancement does not extend to primate cells

To determine whether the enhanced transduction observed with high-concentration FBS and cholesterol was conserved across species, we performed transduction experiments with activated T cells from human and NHP donors. Using titrated GALV-pseudotyped retrovirus, cells were transduced in media containing 10% FBS or 1X cholesterol+30%FBS. In contrast to canine cells, transduction was actually reduced in the presence of high concentration FBS with cholesterol. In human T cells, the negative impact was apparent across all doses, though the difference failed to reach statistical significance (Fig 11A). NHP T cell transduction was quite high under both conditions (>90% at higher virus concentrations) and deviation between the groups was not apparent until 8X dilution when 10% FBS achieved superior transduction. This difference was statistically significant at 32X dilution and beyond (Fig 11B). Taken together, while high concentration FBS and cholesterol enhances transduction in canine cells, this effect does not universally extend to the other species and may adversely affect transduction in primate species.

**Fig 11.**
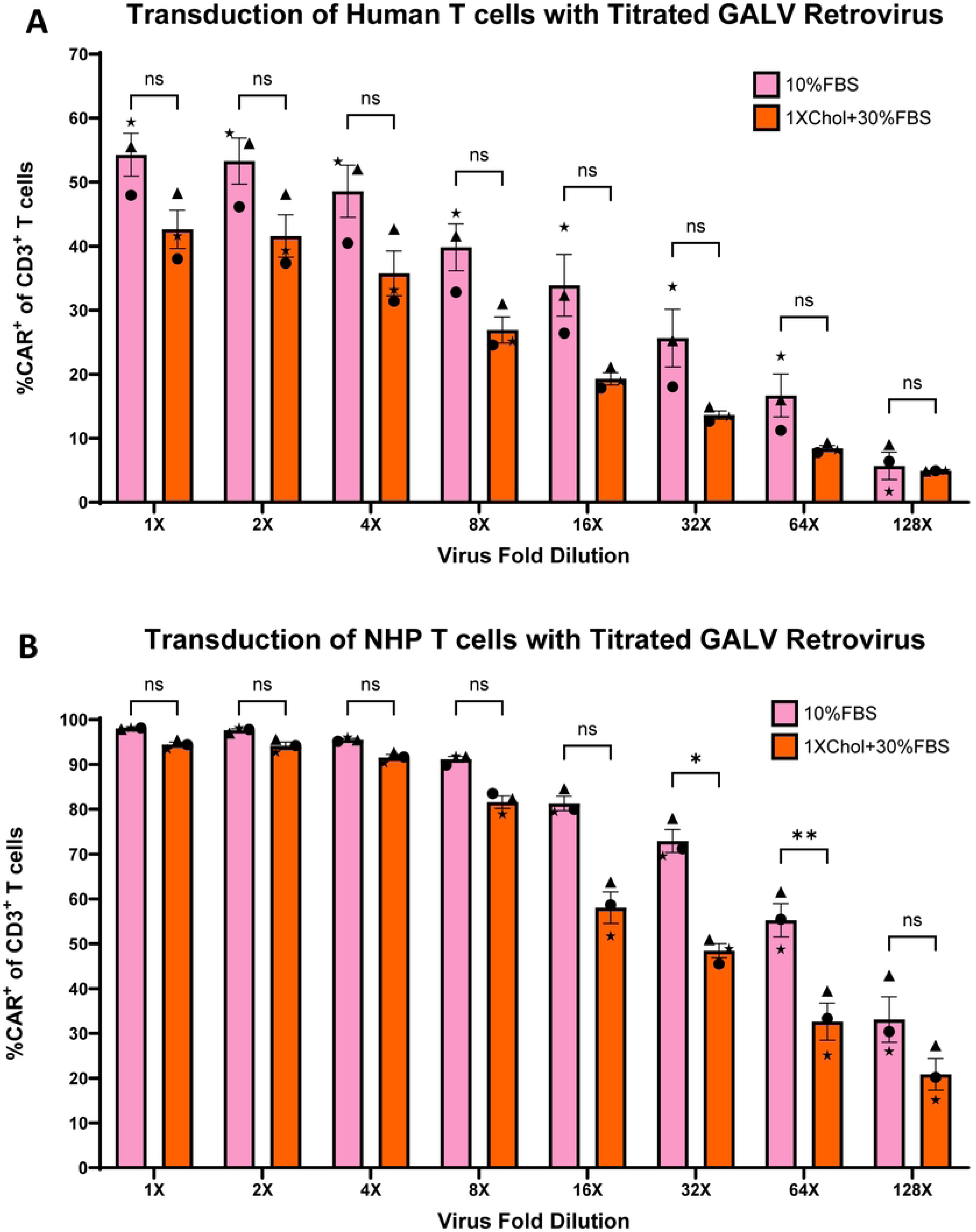
Transduction enhancement using high concentration FBS with cholesterol is not reproducible in primate T cells. After activation, PBMCs from A) three human donors and B) three NHP donors were transduced with GALV-pseudotyped CAR retrovirus in dilutions ranging from 1X (undiluted) to 128X. Transduction was evaluated on day 5 (human) or 3 (NHP) in CD3^+^ T cells. Statistical significance between culture conditions at each virus dilution was assessed by two-way repeated-measures ANOVA followed by Šídák’s multiple-comparisons test. *P < 0.05, **P < 0.01, ***P < 0.001, ****P < 0.0001; ns, not significant. Different symbols represent individual donors.

### Enhanced transduction with FBS/cholesterol is reproducible using autologous canine plasma

We next investigated whether the enhanced transduction observed with FBS and cholesterol could be reproduced using autologous canine plasma. Although FBS can be removed through washing following *ex vivo* CAR T cell expansion, its potential immunogenicity remains a concern, particularly for emerging strategies involving *in vivo* T cell activation and transduction [47–53]. Activated T cell-enriched PBMCs from two canine donors were transduced in media containing 0% or 10% FBS, 1X cholesterol alone, or 1X cholesterol with either 30% FBS or 10% autologous canine plasma (CP). The following day, cells were expanded in media containing 10% FBS and transduction was evaluated after 7 days (S10 Fig). Cholesterol markedly improved transduction efficiency in both donors, with further enhancement when combined with either FBS or CP. Transduction efficiencies with CP were comparable to those achieved with FBS, indicating that autologous CP may serve as a less immunogenic alternative without compromising transduction efficiency.

## Discussion

In this study, we provide a framework for improved canine CAR T cell manufacturing by improving cellular expansion, memory phenotype, and retroviral transduction efficiency. We found that IL-2 supplementation promoted robust expansion and enrichment of less-differentiated memory subsets, and that shorter stimulation periods and increased PBMC seeding density improved transduction efficiency without compromising cellular expansion or memory phenotype. Although implementation of strategies that boost transduction in other species, like phosphate deprivation, did not enhance transduction of canine cells, we discovered that increasing FBS concentrations and cholesterol supplementation substantially improved transduction efficiency. Combining high-concentration FBS with cholesterol further enhanced both transduction and cellular expansion while preserving favorable memory characteristics. This combination also mitigated the reduction in transduction efficiency associated with elevated culture temperatures.

Human clinical studies suggest that CAR T cells with less-differentiated memory phenotypes exhibit greater persistence and are associated with improved patient outcomes [24,54]. Accordingly, we sought to identify cytokine culture conditions that promote T_SCM_ differentiation while limiting the accumulation of terminally differentiated T_EFF_ cells. Sakai et al. [27] previously investigated canine CAR T cell memory differentiation following mitogen-induced stimulation and expansion in IL-2. Marked differences in T_SCM_ and T_EFF_ populations between mitogens suggested that stimulation strategy as well as cytokines play important roles in memory T cell differentiation in dogs. In our previous work, we demonstrated that plate-bound antibody-induced stimulation resulted in better activation, transduction, and expansion than mitogens, while offering a superior safety profile [20]. As cytokines have not been thoroughly evaluated in the context of antibody-induced activation, it became imperative to determine the role of, not only rhIL-2 after antibody stimulation, but of rhIL-7 and rhIL-15 as well.

While human T cell memory subsets are commonly characterized using combinations of CCR7, CD45RA, CD45RO, and CD62L, the limited commercial availability of canine-reactive antibodies has constrained similar analyses in dogs. Previous canine studies have primarily relied on CD62L alone or in combination with CD27 or CD44 [19,26,55,56], limiting the ability to distinguish distinct memory populations. Accordingly, we used CD45RA and CD62L expression to delineate the T cell subsets, enabling a more comprehensive assessment of memory differentiation, particularly T_SCM_ and T_EFF_ populations. We demonstrated that expansion in rhIL-2 supported superior cellular expansion, prolonged activation, and a favorable memory phenotype characterized by increased T_SCM_ and T_CM_ populations and reduced T_EFF_ differentiation. rhIL-7 and rhIL-15 combinations underperformed compared to human literature. This failure may simply be the result of insufficient sequence homology between canine and human IL-7 and/or IL-15 leading to poor binding to canine receptors and thus poor intracellular signaling. It is also possible that these cytokines do not play the same biological role in canine cells as they do in humans. One group defined IL-7 as critical for less differentiated memory subsets in human T cells, but alone failed to drive expansion, which required supplementation with IL-2 or with IL-15 [36]. To fully parse this out, experiments utilizing canine IL-7+IL-15 would be necessary, or exploring a higher range of doses with human IL-7+IL15. However, because the use of rhIL-2 met our specific goals, we did not investigate further.

Increasing initial PBMC seeding density overcame the limited expansion associated with 2-day stimulation while maintaining high transduction efficiency. In initial experiments, stimulation for 2 days resulted in high transduction, but was insufficient for robust expansion. We modified the stimulation conditions slightly by increasing the initial PBMC seeding density in the antibody-coated plates. This simple change overcame the limited expansion previously seen, boosting expansion to ∼100-fold over 7 days. These findings were consistent with human T cell studies that showed higher T cell density improved expansion and viability, attributable in part to the accumulation of secreted catalase, reduced oxidative stress [40], improved paracrine cytokine signaling, and increased opportunity for homotypic T cell synapsing [57,58]. Higher numbers of antigen presenting cells (APCs) in the initial PBMC population may also contribute to some degree [58,59]. However, in the presence of T cell-specific antibody stimulation, resting monocytes and B cells are unlikely to substantially upregulate co-stimulatory molecules that would benefit T cell proliferation.

Phosphate deprivation failed to enhance GALV-mediated transduction in canine T cells, despite previous reports of improved transduction in human and non-human primate lymphocytes [41]. In these studies, phosphate deprivation was proposed to upregulate PiT1 (SLC20A1), a sodium-dependent phosphate transporter that serves as the cellular receptor for GALV entry. Why this approach was unsuccessful in canine T cells remains unclear, particularly given previous evidence that GALV entry in canine cells is mediated by the same receptor [60]. In contrast, we observed in these experiments that serum deprivation significantly reduced transduction efficiency. We further found that transduction improved at high FBS concentrations, demonstrating the importance of serum supplementation in canine T cell engineering. These findings were consistent with human lymphocyte experiments that demonstrated increased transduction with GALV or RD114 pseudotyped virus when serum was present compared to serum free media [46,61,62]. These effects were attributed to the ability of FBS to preserve viral stability over time and also to prevent viral particle aggregation, thereby maintaining infectivity.

Cholesterol/lipid supplementation using a proprietary formula (Gibco 250X Cholesterol Lipid Concentrate) substantially enhanced retroviral transduction efficiency in canine T cells, with further improvements observed when combined with high-concentration FBS (30%). This enhancement extended to both GALV- and RD114-pseudotyped retroviruses, suggesting that the effect is not restricted to a specific viral envelope or receptor-mediated entry pathway. Notably, cholesterol combined with FBS also mitigated temperature-associated reductions in transduction efficiency, potentially reflecting improved viral stability or altered cellular processes that facilitate transduction. Beyond transduction, this combination nearly doubled T cell expansion while preserving less differentiated memory T cells and avoiding T_EFF_ terminal differentiation, suggesting that these culture conditions may improve both the efficiency and quality of canine CAR T cell manufacturing. The lack of enhancement with BSA suggests that other components of FBS may be responsible for the synergistic effects observed with cholesterol supplementation.

We propose four possible mechanisms for enhanced transduction rates with cholesterol and high concentration FBS. 1) Enhanced viral stability. We demonstrated FBS and cholesterol supplementation largely prevented temperature-associated reductions in transduction efficiency. The combination of FBS and cholesterol may cooperatively enhance viral particle stability, an effect previously proposed for FBS supplementation alone [46], but not cholesterol alone [63]. 2) Increased mitotic activity. The combination of FBS and cholesterol substantially increased T cell proliferation, potentially providing more opportunities for retroviral integration. Since gammaretroviral transduction depends on mitotic nuclear envelope breakdown for access to host DNA, increased mitotic activity may contribute to enhanced transduction efficiency [39]. 3) Increased lipid raft formation. Cell receptors that viruses exploit to gain entry into the cell are often contained in lipid rafts in the cellular membrane. It is possible that cholesterol/lipid supplementation drives upregulation of cholesterol synthesis, thus increasing lipid raft formation [45,64]. 4) Improved viral endocytosis and endosomal escape. It is possible that cholesterol supplementation prepares a cell membrane environment that better facilitates viral entry and endosomal escape for enhanced access to cytoplasm [64–66]. Importantly, the improved transduction seen in canine cells was not reproducible in human and NHP cells. Prior work reported that cholesterol supplementation during viral production improved retroviral and lentiviral transduction, but that same study saw no benefit in mouse fibroblasts when cholesterol was added during transduction [63]. This species variance may provide insight and guide future investigations into possible mechanisms.

There are several limitations to the current study. We recognize that T cell toxicity may be an issue. Our approach seems to promote uptake of virus particles, even with low-titer virus. High multiplicities of infection (MOI) increase the likelihood of multiple vector integrations per cell. This can increase cellular stress and the risk of genomic disruption, resulting in the increased potential for cell death or neoplastic transformation [65–67]. Determining optimal cholesterol and FBS dosing with titrated virus will be important for researchers adopting this protocol to reduce the risk of these issues. Also, we limited experiments to GALV and RD114-pseudotyped gammaretrovirus. We do not know if similar enhancement can be demonstrated with other retroviral envelopes described for canine transduction such as VSV-G or 10A1 [26,60]. It is also unclear if this strategy will enhance lentiviral-mediated transduction [19].

In summary, we report methods to improve canine CAR T cell production, characterized by high transduction efficiency with maintained CAR expression, excellent expansion capacity with preserved T_SCM_ phenotypes, and excellent viability that will likely confer effective and enduring anti-tumor activity. Additionally, we present a novel way to boost transduction efficiency by transducing T cells in the presence of cholesterol and high concentration FBS. This offers substantial improvement in transduction, while avoiding repeated rounds of viral transduction or co-culture with target cells. In several experiments with multiple donors, transduction rates were consistently above 70-80%, with the highest rate of 92.0% achieved. In fact, we demonstrate with our method that 16-fold less virus is needed to achieve the same transduction efficiency attained with a standard transduction protocol. This, along with the enhanced CAR T cell expansion resulting from our methods, translates to reduced culture time, faster patient treatment, and substantial cost savings. It also represents a CAR T cell engineering protocol that is highly accessible to researchers and clinicians. We hope our findings will make a significant impact in the field of canine CAR T cell engineering and will hasten CAR T cell immunotherapy as a standard of care for cancer treatment in dogs.

## Acknowledgements

We are grateful to Dr. Nicola Mason (UPenn) for her bountiful knowledge, guidance, and for facilitating essential collaborations. Thank you to Dr. Peter Moore and Kristy Harmon (UC Davis) for assistance in procuring the canine specific antibodies used in this study; Brian Hayes (Fred Hutchinson Cancer Center) for gifting the 5B8 hybridoma cell line; Dr. Gianpietro Dotti (UNC) for gifting FLYRD18 cells and a stably transduced FLYRD18 gammaretroviral GFP producer cell line. Additionally, we thank NCSU Laboratory Animal Resources veterinary medical team for providing the canine whole blood essential for our experiments. Lastly, we thank Ramiro Diz, Bob Immormino, and Ayrianna Hedgecock (UNC Flow Core) for their assistance with flow sorting and with flow cytometry data collection, troubleshooting, and analysis. This work was supported by the National Institutes of Health through Grant Award R37-CA260223 from the NCI, seed funding from the Comparative Medicine Institute and the Lineberger Cancer Center and by start-up funds provided by the University of North Carolina at Chapel Hill, North Carolina State University at Raleigh, and the Lineberger Cancer Center.

## Supporting information

**S1 Fig. Gating strategy for determining memory T cell subsets.** A representative plot to show memory subset gating strategy. Lymphocytes are gated in SSC/FSC plot. After exclusion of doublets, viability was determined by the exclusion of cells positive for LIVE/DEAD Near IR viability dye. T cells were identified by gating on CD5^+^ cells. Finally, CD62L and CD45RA expression were evaluated. Phenotype was determined according to quadrant: Upper right quadrant – Naive/stem cell memory (T_SCM_) T cells; upper left – central memory (T_CM_) T cells; lower left – effector memory (T_EM_) T cells; and lower right – effector (T_EFF_) T cells.

**S2 Fig. Canine T cell activation occurs early in the stimulation period.** PBMCs from four canine donors were stimulated with plate-bound antibodies and activation was evaluated on days 1-5 as determined by cell surface expression of CD25 on CD5^+^ T cells (AB=antibody stimulated). Horizontal lines indicate mean values, and error bars represent standard error of the mean. Different symbols represent individual donors.

**S3 Fig. Stimulation of canine T cells for 2 days favors higher CD4 proportionality, while 2- and 3-day stimulation promotes less differentiated memory subsets.** PBMCs from four canine donors were activated with stimulatory antibodies for 2-5 days. After removal from stimulation, T cell-enriched PBMCs were transduced. Non-transduced cells (NT) were stimulated for 5 days, but were not transduced. A) On day 4 of expansion, CD4/CD8 subsets were determined in the CD5^+^ T cell population (DN=CD4^-^CD8^-^ double negative, DP=CD4^+^CD8^+^ double positive) and B) the proportion of CD4^+^ T cells in particular. C) Also, memory phenotypes were determined and D) the proportion of T_SCM_ cells in particular. For B) and D), horizontal lines indicate mean values, and error bars represent standard error of the mean. Pairwise statistical analysis was performed by one-way ANOVA with Tukey’s multiple means comparison; * p<0.05, ** p<0.01, ns not significant. Different symbols represent individual donors.

**S4 Fig. Gating strategy for determining CD4/CD8 T cell subsets.** A representative plot to show CD4/CD8 subset gating strategy. Lymphocytes are gated in SSC/FSC plot. After exclusion of doublets, viability was determined by the exclusion of cells positive for LIVE/DEAD Near IR viability dye. T cells were identified by gating on CD5^+^ cells. Finally, CD4 and CD8 expression were evaluated and phenotype was determined according to quadrant: Upper left quadrant – CD4^+^ cells; upper right – CD4^+^CD8^+^ double positive (DP) cells; lower left – CD4^-^CD8^-^ double negative (DN) cells; and lower right – CD8^+^ cells.

**S5 Fig. Phosphate deprivation prior to transduction does not improve transduction efficiency with GALV-pseudotyped retrovirus in canine T cells.** A) Schematic of the experiment design. PBMCs from four canine donors were activated with stimulatory antibodies for 2 days. On the day of transduction, standard 10% FBS T cell media was replaced with 3% FBS, phosphate free media 16 and 6 hours before transduction. The “No Deprivation” group was maintained in standard media. T cell-enriched PBMCs were returned to standard media at the time of transduction and B) transduction efficiency was evaluated for CD5^+^ T cells on days 4 and 7. Horizontal lines indicate mean values, and error bars represent standard error of the mean. Statistical analysis was performed by two-way ANOVA with Tukey’s multiple means comparison; * p<0.05, ** p<0.01, *** p<0.001, **** p<0.0001, ns not significant. Different symbols represent individual donors.

**S6 Fig. Enhanced canine T cell transduction with high concentration FBS and cholesterol is confirmed in a repeat experiment.** PBMCs from four canine donors were activated with stimulatory antibodies, then transduced with GALV pseudotyped CAR retrovirus. During the transduction period, standard media was replaced with media containing 10% FBS, 1X cholesterol, or 30% FBS with cholesterol. The following day, cells were transferred to standard 10% FBS media and expanded. Transduction was evaluated on day 7. Horizontal lines indicate mean values, and error bars represent standard error of the mean. Pairwise statistical analysis was performed by One-way ANOVA with Tukey’s multiple means comparison; ** p<0.01, ns not significant. Different symbols represent individual donors.

**S7 Fig. Gating strategy for determining transduced T cells.** A representative plot to show transduced T cell gating strategy. Lymphocytes are gated in SSC/FSC plot. After exclusion of doublets, viability was determined by the exclusion of cells positive for LIVE/DEAD Near IR viability dye. T cells were identified by gating on CD5^+^ cells. Finally, transduced cells were identified by labeling the G4S linker on the extracellular portion of the CAR protein.

**S8 Fig. Enhanced expansion with FBS/cholesterol does not drive terminal differentiation of memory T cells.** Activated PBMCs from three canine donors were transduced with GALV-pseudotyped CAR retrovirus. During the transduction period, cells were either maintained in standard 10% FBS T cell media or transduced in media containing 30% FBS with different concentrations of cholesterol. The following day, cells were transferred to standard 10%FBS for expansion. On day 4, memory T cell subsets were determined including A) The proportion of T_SCM_ + T_CM_ cells, B) the proportion of T_CM_ cells alone, C) the proportion of T_SCM_ cells alone, and D) the proportion of terminally differentiated T_EFF_ cells. Horizontal lines indicate mean values, and error bars represent standard error of the mean. Pairwise statistical analysis was performed by one-way ANOVA with Tukey’s multiple means comparison; * p<0.05, ns not significant. Different symbols represent individual donors.

**S9 Fig. High-concentration FBS and cholesterol mitigate temperature-associated reductions in transduction efficiency with reduced donor variability.** PBMCs from four canine donors were activated with stimulatory antibodies, then transduced with GALV-pseudotyped CAR retrovirus. During the transduction period, cells were either maintained in standard 10% FBS T cell media or transferred to media containing 1X cholesterol, 30% FBS (+/- cholesterol), or BSA (+/- cholesterol). Cells were cultured at 37°C or 38.8°C overnight. The next day, cells were transferred to standard media with IL-2 and expanded at 38.8°C. Changes in transduction were evaluated under these different temperature culture conditions. A) Relative reduction (%) for each donor was evaluated between groups. Pairwise statistical analysis was performed by one-way ANOVA with Tukey’s multiple means comparison; * p<0.05, ns not significant. Different symbols represent individual donors. B) The average transduction at lower and higher temperatures was directly compared. Statistical significance was assessed by two-way repeated-measures ANOVA followed by Šídák’s multiple-comparisons test. ***P < 0.001.

**S10 Fig. Autologous canine plasma with cholesterol replicates the transduction-enhancing effects seen with FBS.** Activated T cell-enriched PBMCs from two canine donors were transduced with GALV-pseudotyped retrovirus in media containing 10% FBS or 1X cholesterol with or without 30% FBS or 10% autologous canine plasma (CP). The following day, cells were expanded in media containing 10% FBS and transduction efficiency in CD5^+^ T cells was evaluated on day 7 post-transduction.

**S1 Table. Donor information.** A summary of pertinent medical information for the seven canine blood donors used in this study.

